# Molecular kinetics dictate population dynamics in CRISPR-based plasmid defense

**DOI:** 10.1101/2025.03.27.645723

**Authors:** Luke Richards, Danna Lee, Jakub Wiktor, Axel Truedson, Johanna Cederblad, Daniel Jones

**Affiliations:** Department of Cell and Molecular Biology, Uppsala University, Husargatan 3 Box 596, 75124, Uppsala, Sweden

**Keywords:** CRISPR, plasmid conjugation, population dynamics, microfluidics, single-cell tracking

## Abstract

Understanding and manipulating mobile genetic element (MGE) spread within bacterial communities represents a great challenge with wide-ranging potential benefits. It requires a detailed understanding of spatial effects and cell-to-cell variability in defence systems such as CRISPR-Cas. Here, we report a time-lapse imaging-based assay with simultaneous fluorescent labeling of CRISPR-Cascade complexes and the conjugative plasmid RP4, enabling direct, single-cell-resolution observation of conjugation and CRISPR interference. We find that Cascade, under wild-type-like expression scenarios, provides a nudge towards plasmid clearance rather than full immunity, and can be counteracted by the ParDE plasmid addiction system. These limitations could be overcome by higher Cascade expression, yet by quantifying single-cell variability, we observe a tradeoff between plasmid clearance and cell growth. We synthesize these measurements into a spatially-resolved, agent-based model of plasmid population dynamics. The imaging and analysis techniques used here will facilitate disentanglement of how single-cell molecular events result in community-wide plasmid dynamics.

## 1 Introduction

Microbial communities are ubiquitous and extremely complex, taxonomically, spatially and functionally [1, 2]. They possess an added level of complexity driven by their propensity to share genetic material (and associated functions) via horizontal gene transfer (HGT). Plasmid-mediated HGT, often transferred via Type IV secretion systems (T4SS) [3], can bestow beneficial functions on plasmid hosts. Plasmids can encode, for example, new metabolic functions [4, 5] or antibiotic resistance. The latter is driving a burgeoning public health crisis [6]. Despite their host-benefiting features, conjugative plasmids also impose a fitness cost [7–10]. Bacteria have consequently evolved an array of defense systems against conjugative plasmids and broader mobile genetic elements (MGEs) [11]. Equally, those mobile elements have evolved strategies to evade such defense systems [12]. Plasmid acquisition and maintenance cost must be balanced against defense cost and the current - or indeed potential future - benefit of the encoded functions.

CRISPR defense is widely recognized as a method for microbes to protect against invading genetic material [13, 14] with CRISPR systems present in around 75% of archeal isolates and around 36% of bacteria [15]. Nonetheless, the evidence for CRISPR-Cas systems’ effectiveness is mixed [16, 17]. In general, CRISPR appears to be used by bacteria to target plasmids for degradation, although perhaps less often than phages [18]. Additionally, genetic studies have demonstrated a negative correlation between CRISPR defense, and plasmids, phages and multi-drug resistance [19, 20]. Despite CRISPR’ s apparent targeting of plasmids, experiments with *E. coli* have demonstrated plasmids persisting over many cell generations whilst being targeted by CRISPR [21]. By extension, mere CRISPR presence is not a panacea against plasmid invasion. CRISPR-based defense variation within populations could be a “bet-hedging” strategy to maintain low plasmid frequency. This has been discussed as a method for dealing with environmental uncertainty [22].

Wild-type *E. coli* K-12 harbors a Type I-E CRISPR-Cas system, comprised of the multi-protein Cascade complex for target search, which subsequently recruits Cas3 helicase-nuclease for degradation. *E. coli* K-12 generates very low levels of CRISPR-RNA, suggesting that this system is perhaps not active [23] under known environmental conditions. Indeed, in the case of *E. coli*, CRISPR interference seems ineffective against mobile genetic elements [24]. Nonetheless, the system’ s components are fully functional [23], giving this model organism an important role in understanding CRISPR-MGE dynamics. *E. coli* studies artificially inducing CRISPR components show a clear relationship between Cascade expression and plasmid clearance and estimate a 90 minute search time per Cascade complex [25]. Single-cell Cascade expression stochasticity demonstrably affects CRISPR-targeting efficacy [26] but highly-expressed CRISPR systems carry costs in terms of expression burden, DNA damage and replication interference [27, 28]. Furthermore, growth rate of cells in and of itself has been implicated in cells’ ability to degrade plasmids [26]. Taken together, these studies indicate a complex interplay between defense expression growth, clearance, and cost-benefit trade-offs at a single cell level, which play out to become population level changes.

Theoretical studies of plasmid conjugation and population dynamics often take the form of “mass action” ordinary differential equation (ODE) models which describe well-mixed bacteria in homogeneous environments [29–31]. However, real-life bacterial habitats are characterized by spatial constraints, physical inhomogeneity, small population sizes, and cell-to-cell variability not captured by ODEs. Confined environments can affect conjugation dynamics [32, 33] and cell-to-cell variability may have adaptive potential [26]. Understanding such factors is, therefore, key to describing population dynamics.

We investigate the population and single-cell dynamics of plasmid conjugation and CRISPR interference in *E. coli* using microfluidics-coupled, time-lapse fluorescence microscopy. These methods facilitate measurement of key biophysical parameters in the conjugation - CRISPR dynamic. At the population level we find resistance to conjugative plasmid spread when CRISPR interference is high. This also has profound consequences for the population depending on the presence of a plasmid addiction system. With the help of single-cell tracking and classification, we are able to report, for the first time, conjugation rates per neighboring plasmid-containing cell and estimate the latent period between plasmid acquisition and subsequent onward transmission. We also find that cell-to-cell variability in CRISPR-Cascade expression significantly affects dynamics at the single-cell level, inducing a growth cost at high expression levels where the clearance rate is highest. The insights presented here advance our understanding of conjugation dynamics in natural and synthetic microbial communities and build towards potential future manipulation across medical, industrial and agricultural applications.

## 2 Results

### 2.1 Population dynamics across CRISPR interference levels

To visualize plasmid population dynamics we engineered ‘donor’ and ‘recipient’ *E. coli* cells such that donor cells displayed no fluorescent properties while recipient cells displayed constitutive, homogeneous mCherry-ParB expression, and inducible Cas3 and Cascade operon expression (by IPTG and arabinose, respectively). Cas8e, part of the Cascade complex, was translationally fused to sYFP2, and sYFP2-Cas8e was used as a proxy for Cascade concentration throughout this work [25]. Transconjugant cells therefore also exhibit sYFP2-Cascade expression but form fluorescent mCherry foci from mCherry-ParB binding to *parSMT1* sites on RP4 (Fig. 1A). We observed conjugation and Cascade expression dynamics in microfluidic chips (Supplementary Fig. S1) over 6 hours with 6 RP4 plasmid variants; containing either 0, 1 or 18 spacer targets, each with and without *parE* from RP4’ s *parDE* toxin-antitoxin system. See Methods for more details on plasmids and strains.

**Fig. 1.**
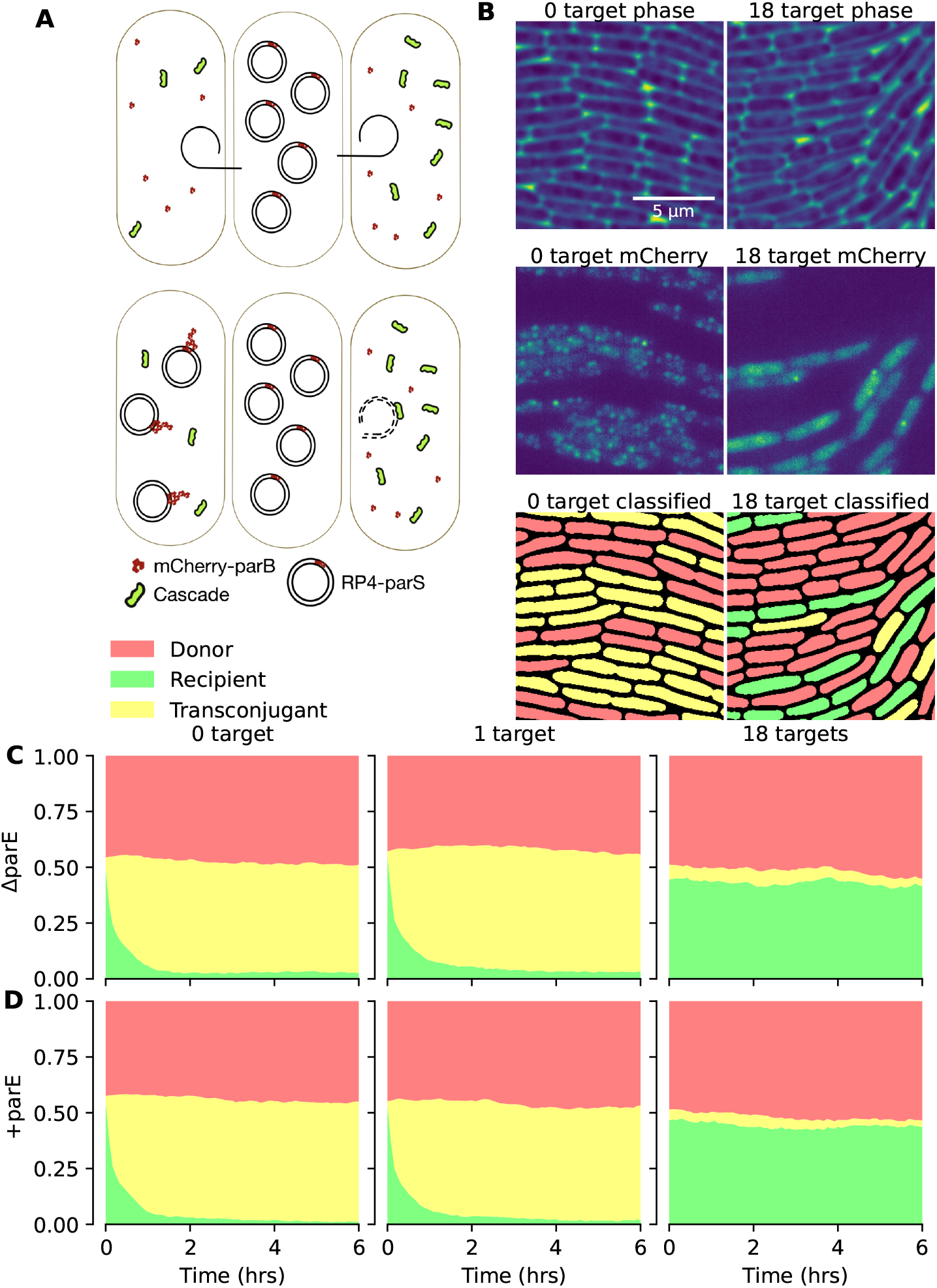
A) Donor cells harboring parSMT1-containing RP4 plasmids transfer plasmids via conjugation into fluorescently labeled recipient cells. Recipient cells contain mCherry-tagged ParB and sYFP2-tagged Cascade to allow tracking of conjugation and Cascade expression levels. B) Corresponding phase contrast (top), mCherry fluorescence (middle), and classified segmented cell (bottom) images for 0 target and 18 target RP4 variant experiments. C) Mean fraction of the cell population classified as donor, recipient or transconjugant across the course of microfluidic experiments with RP4 plasmid variants. At least 8000 individual cells were observed for each time point.

To start experiments, a 50/50 mixed population of donor and recipient cells was loaded into the microfluidic device and time-lapse imaging was performed in phase contrast, mCherry, and sYFP2 channels for 6 hours. Observing the population fraction corresponding to each cell class (donor, recipient, or transconjugant) presents an overview of population dynamics over time for each CRISPR interference level (Fig. 1CD). For the non-targeted RP4-0-target variants, the recipient population is rapidly and comprehensively converted to transconjugants, reflecting RP4’ s high conjugation rate. For the RP4-1-target variants, the transconjugant takeover is slightly slowed but broadly similar to the 0-target case. Finally, the RP4-18-target variants are efficiently targeted by CRISPR, with only a very small proportion of the population succumbing to conjugation.

The presence or absence of *parE* did not substantively alter population dynamics in this instance (Fig. 1), somewhat surprisingly as cleared cells should experience ParE toxicity. However, in these experiments Cascade induction took place prior to the experiment; conjugation is never allowed to proceed in the absence of CRISPR defense. We hypothesized that, under these conditions, the toxin may not have time to accumulate inside cells before clearance occurs.

#### 2.1.1 Toxin-antitoxin systems can be effective counter-defense against CRISPR interference

To assess the impact of plasmid establishment in cells before CRISPR interference is induced, RP4-18-target donor cells were introduced into microfluidic experiments mixed with non-induced recipient cells. For the first hour of imaging, conjugation was allowed to proceed in the absence of CRISPR induction, at which point media was switched to media containing inducers (Fig. 2). Around the 70-80 minute time point the clearing of transconjugants is apparent in both the +*parE* (right) and Δ*parE* (left) experiments, as well as a concomitant increase in Cascade expression (Fig. 2B). Additionally in the case of +*parE* there is also a marked decrease in the fraction of the population made up of recipients (Fig. 2A). In Fig. 2C we show representative images of segmented and classified cells at the end of the experiments. Experiments with *parE* present on RP4 show elongated, dying recipient (cleared transconjugant) cells whereas without *parE* the cells clear and remain healthy in appearance. Control experiments were also performed where the recipient cells had CRISPR arrays deleted. In these control experiments cells were never cleared of plasmids (Fig. S2). Taken together, these observations suggest that CRISPR interference may be an ineffective mechanism to protect against unwanted conjugative plasmids that make use of *parDE*, and potentially other toxin-antitoxin, systems. The results also highlight the importance of timing of MGE defence system expression [34], insofar as CRISPR-Cas expression after plasmid establishment came too late to mitigate TA toxicity.

**Fig. 2.**
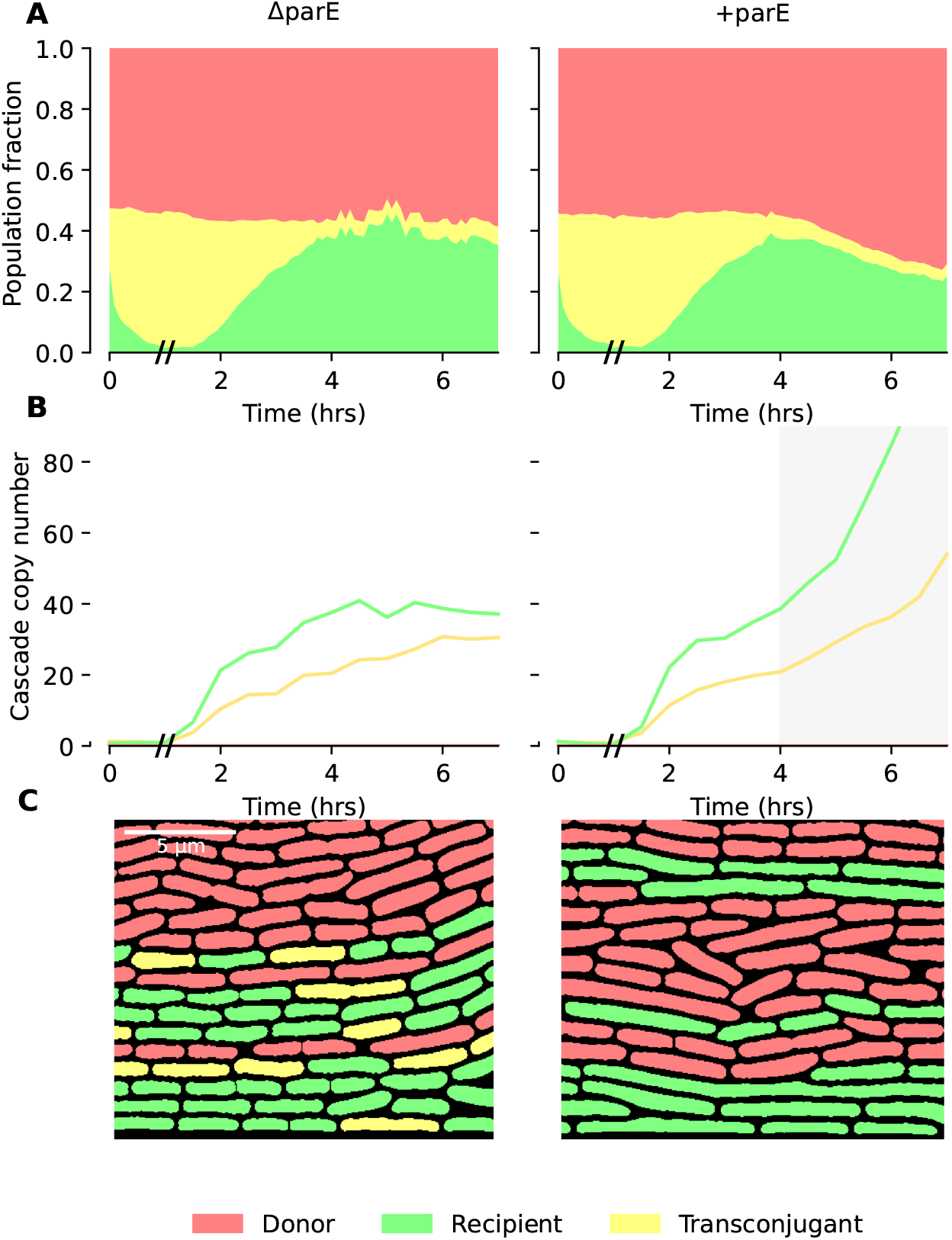
A) Mean fraction of the cell population classified as donor, recipient or transconjugant across the course of microfluidic experiments with delayed Cascade induction with 18-target RP4 plasmids, with and without *parE*. B) Median Cascade copy number per cell across the course of the experiment. Inducers were added to the media at 60 minutes, indicated with // on axis. In the case of +*parE*, copy number estimation beyond around 4 hrs becomes unreliable as dying cells become very bright in the sYFP2 channel. C) Representative segmented, classified cell images at the end of +*parE* (right) and Δ*parE* (left) experiments. At least 5000 individual cells were observed for each time point.

### 2.2 Single-cell characterization of conjugation kinetics

#### 2.2.1 Conjugation proceeds with a single rate-limiting step at a rate of two conjugation events/hour/donor neighbor

The population dynamics shown in Figs. 1 and 2 reflect the interplay of the opposing forces of conjugation and CRISPR interference. Next, we sought to characterize each of these processes separately at the single-cell level, focusing first on conjugation. In Fig. 3, we examine how quickly recipient cells are “converted” to transconjugants as a function of how many donor neighbors they have. In particular, in Fig. 3A, we plot the relative transconjugant fraction *T/*(*T* + *R*), where *T* is the number of transconjugants and *R* is the number of recipients. This quantity represents the fraction of recipients that have been converted to tranconjugants. We plot this fraction for different subsets of the population based on how many donor neighbors each recipient or transconjugant has (see Methods, “Image Analysis”). Recipients with more donor neighbors are conjugated more rapidly than recipients with fewer donor neighbors.

**Fig. 3.**
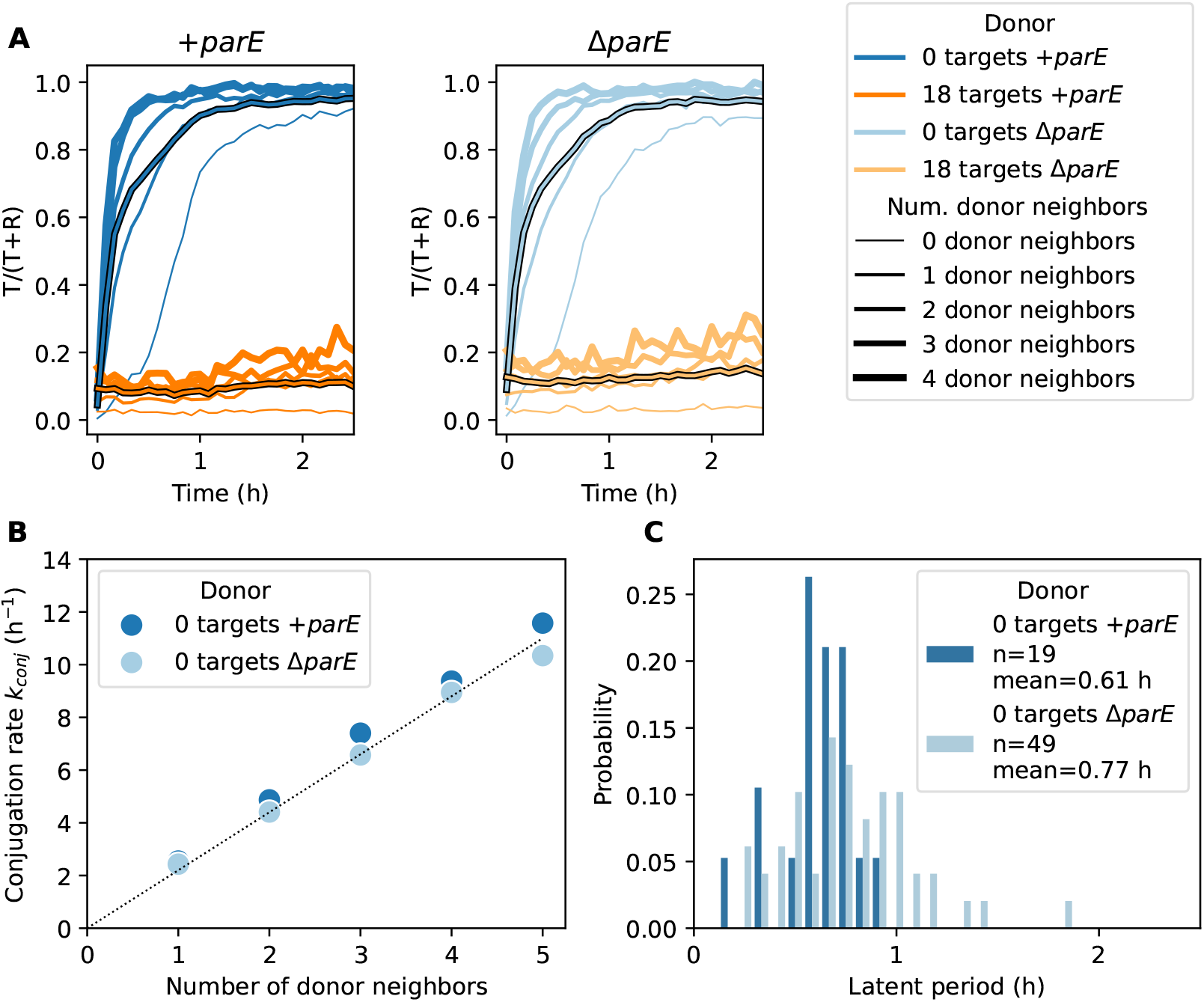
Single-cell characterization of conjugation dynamics. **(A)** Relative transconjugant fraction *T/*(*T* + *R*) as a function of time, donor plasmid (color) and number of donor neighbors (line thicknesses). *T* and *R* denote numbers of transconjugants and recipients, respectively. Black-outlined lines are averages across all donor-neighbor counts. CRISPR targeting makes recipients much less likely to become transconjugants, but cells with more donor neighbors are more likely to become transconjugants. At least 2184 cells were analyzed for each donor and timepoint (3182 cells on average). 1-target plasmids have similar dynamics to 0-target plasmids (omitted for clarity). **(B)** Dependence of conjugation rate on number of donor neighbors. Conjugation rates were determined by fitting exponential functions (Equation 1) to the 0 target donor curves shown in (A) for nonzero donor neighbors. The conjugation rate is directly proportional to the number of donor neighbors, highlighting the effects of local spatial inhomogeneities. Dotted line is a guide to the eye. **(C)** Latent period for conjugation, defined as the length of time from when a recipient becomes a transconjugant until the new transconjugant itself performs conjugation (see text).

To isolate the dynamics of conjugation, we further analyzed RP4-0-target donors which are not targeted for CRISPR interference. For each of the curves with one or more donor neighbors, we fit an exponential function of the form

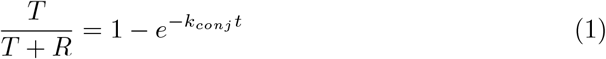

where *k*_*conj*_ represent the conjugation rate for a given number of donor neighbors. In Fig. 3B, we plot *k*_*conj*_ for donor neighbor counts from 1 to 5 and find that conjugation rate is linearly proportional to donor neighbor count. Thus, conjugation from donor neighbors appears to be independent and additive. For single donor neighbors, we obtained a value of *k*_*conj*_ = 2.4 h^−1^, which implies that recipients with a single donor neighbor will be conjugated after 1*/*2.4 h^−1^ *≈* 0.42 h or 25 minutes on average. The mean numbers of donor neighbors during the first hour of the experiment were 2.0 and 1.9 for +*parE* and Δ*parE* RP4-0-target donors, respectively.

#### 2.2.2 New transconjugants experience a 40-50 minute long latent period before performing conjugation

In Fig. 3A we also plotted *T/*(*T* + *R*) for cells with zero donor neighbors. These curves generally start to rise after a delay of approximately 30 minutes from *t* = 0. We surmised that this delay reflects the latent period required for newly-conjugated transconjugants to produce conjugation machinery and become able to perform conjugation themselves. However, as these plots are based on the number of donor-neighbors in a particular frame only, some of these cells may have transiently come into contact with donors in previous frames, meaning that 30 minutes should be considered a lower bound for the latent period.

To more accurately characterize the latent period, we turned to lineage-based analysis based on tracking individual cells and their daughters over time [35]. We sought to identify lineages of recipient cells in which conjugation is performed by newly-conjugated transconjugant cells rather than donor cells (see Methods for details) In Fig. 3C, we plot the times from the first appearance of a transconjugant neighbor until the recipient is conjugated and find a mean latent period of approximately 40-50 minutes, roughly consistent with the picture in Fig. 3A.

#### 2.2.3 Cell-to-cell variability in Cascade expression determines clearance rates

To investigate the function of expression stochasticity in potential bet-hedging strategies we now turn to analyzing the variability in Cascade expression between single cells (Fig. 4A). In Fig. 4B, we plot a histogram of Cascade copy number per cell for recipients and transconjugants, combined from all experiments and donor strains. On average, recipient or transconjugants contained 42 Cascade complexes per cell. With single-cell Cascade copy number estimations in hand, we evaluated the extent to which Cascade expression is costly for cell growth. In Fig. 4B, we also plot the growth rate as a function of binned Cascade expression (blue dots; see right y-axis). We find that growth rate declines roughly linearly with Cascade expression, decreasing by about 5% at 150 Cascade per cell. This suggests a non-negligible tradeoff between growth and defence.

**Fig. 4.**
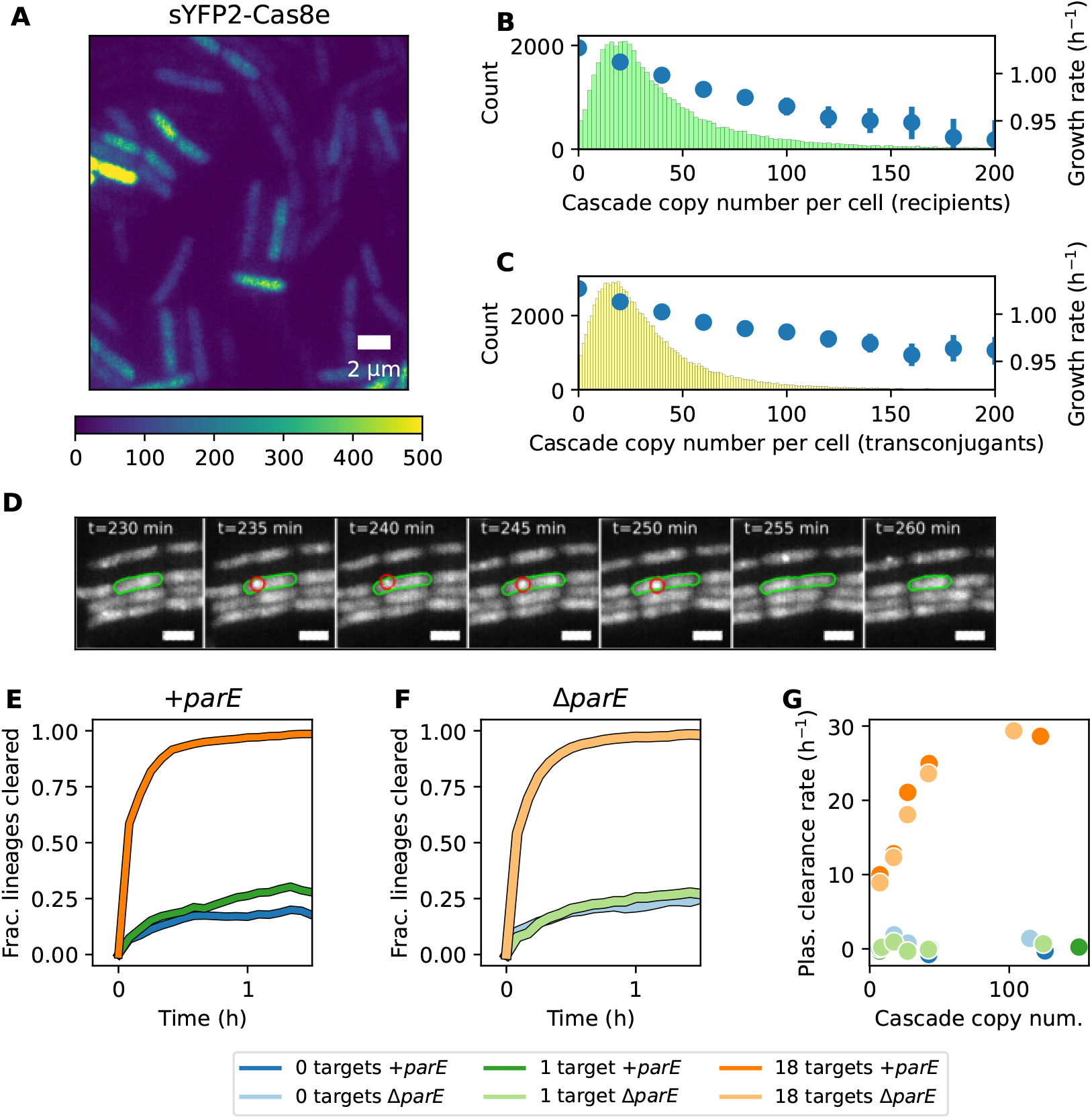
Cell-to-cell variability in CRISPR interference. **(A)** Representative fluorescence image acquired in the Venus channel. The image has been rescaled so that the colormap indicates the corresponding Cascade copy number in an average-sized cell. Dark areas correspond to donor cells, which do not express sYFP2-Cas8e. **(B)** and **(C)**. Estimated Cascade copy number per cell and dependence of growth rate on (binned) Cascade copy number for recipient and transconjugant cells. **(D)** Time-lapse images in mCherry-ParB channel showing a recipient cell (green outline) that receives a plasmid (dot outlined in red) at *t* = 235 min before clearing the plasmid at *t* = 255 min. **(E)** and **(F)**. Fraction of cell lineages that acquired and subsequently cleared the plasmid as a function of time and CRISPR target number (color). Initial plasmid acquisition is defined as *t* = 0 in these plots. More CRISPR targets and higher Cascade expression generally yield faster clearance. At least 1189 unique plasmid infection events were analyzed for each donor. **(G)** Plasmid clearance rate for all plasmid variants as a function of Cascade expression. Clearance rate increases linearly for 18-target plasmid before saturating at high Cascade concentration. Limited clearance observed for 1-target plasmid. See text and Appendix A for calculation of clearance rate.

Next, we turned our attention to the dynamics of CRISPR interference. We performed cell tracking to build a database of “infection events” - *i*.*e*. intervals of a cell lineage during which plasmids are present, illustrated in Fig. 4D. By summing over infection events, we can compute the fraction of lineages that have cleared the plasmid as a function of the elapsed time since initial conjugation. Note that this can involve multiple generations of cells along a lineage. This clearance fraction is shown in Figs. 4E and F for *+parE* and Δ*parE* plasmid variants, respectively. Considering plasmid variants first, we find rapid and complete clearance for RP4-18-target variants, an extremely low level of clearance for RP4-1-target variants, and low albeit nonzero clearance for RP4-0-target variants. The latter should have essentially zero clearance and is therefore representative of a false negative dot-detection rate (plasmid loss due to cell division may also contribute). After about 20-30 minutes, the RP4-0-target clearance curves plateau around 0.2, indicating that plasmid copy number has built up sufficiently that the probability of false negatives is small.

Considering Cascade expression variability, we binned “infection events” into quintiles according to mean Cascade expression during the infection. We then estimated the plasmid clearance rates for each plasmid variant and Cascade expression quintile. To do so, we used the clearance fraction at *t* = 5 min after initial infection. Assuming that plasmid replication time is greater than 5 min, this allows us to avoid the confounding effects of plasmid replication after conjugation. We further corrected for false negative plasmid measurements and the possibility of additional conjugation events during the 5 minute interval; see Appendix A for a complete derivation. The estimated clearance rates are shown in Fig. 4G as a function of Cascade copy number, and are also reproduced in Supplementary Table S1. Consistent with Fig. 4EF, the clearance rate for 0-target and 1-target plasmids are close to zero, while 18-target clearance rates increase linearly with Cascade before plateauing around 50-100 Cascade/cell. For Δ*parE* RP4-18-target, the estimated clearance rate was 18 h^−1^ for the 27 Cascade/cell bin, yielding a per-Cascade clearance time of (18 h^−1^)^−1^ *×* 27 = 1.5 h. In the Discussion we compare these rates to previous biophysical measurements and find good agreement [25].

### 2.3 Bioinformatics: ‘Wild’ toxin-antitoxin and CRISPR spacer relationship

Given our earlier finding that, in the absence of strong prior induction of Cascade, *parE* -containing plasmids are able to effectively counter CRISPR clearance and cause cell death, we hypothesized that spacers against *parE* -containing plasmids should be underrepresented in CRISPR arrays found in bacterial populations. Effective CRISPR targeting and *parE* should be evolutionarily incompatible. A negative correlation between the presence of *parE* and CRISPR targets in available sequence databases would provide strong evidence for the applicability of this mechanism in wider bacterial ecology.

We searched 72556 plasmid sequences [36] for spacers and *parE* using 11767782 known spacer sequences [37] and *E. coli* RP4 *parE* (XCD26522.1). Consequently we identified 42728 plasmids with spacer target sequences, 502 plasmid sequences containing *parE* -like sequences and 106 sequences with both *parE* and CRISPR spacer targets. Of the spacer-targeted plasmids, the maximum and minimum number of spacers was 288 and 1. The mean and median numbers were 16.94 and 6, respectively. If all plasmids are considered those values drop to 9.97 and 1. Naïvely, the global fraction of spacer-targeted plasmids 42728/72556 (59%) compared to the fraction of spacer-targeted, *parE* -containing plasmids 106/502 (21%), is an under-representation of spacer targeting in *parE* -containing plasmids.

To test this relationship statistically we constrained our analysis to ten host species where *parE* -containing plasmid presence was most likely and constructed a phylogeny-weighted random permutation test (4). Plasmid phylogenies were constructed using gene presence-absence (4). Qualitatively, we can see that the presence of both *parE* and CRISPR is not restricted to any one species or phylogenetic lineage but bias is apparent (Fig. 5). Co-occurrence of *parE* and CRISPR targets is observed less frequently than expected by chance, when accounting for phylogeny (Fig. 5 and Table S5). We repeated the method for eight other known toxin genes. We found that *relE, vapC* and *ccdB* actually showed more co-occurrence than expected by chance (table S5), suggesting strong CRISPR targeting in these cases. The effectiveness of such targeting against toxin-antitoxin systems is likely highly context-dependent.

**Fig. 5.**
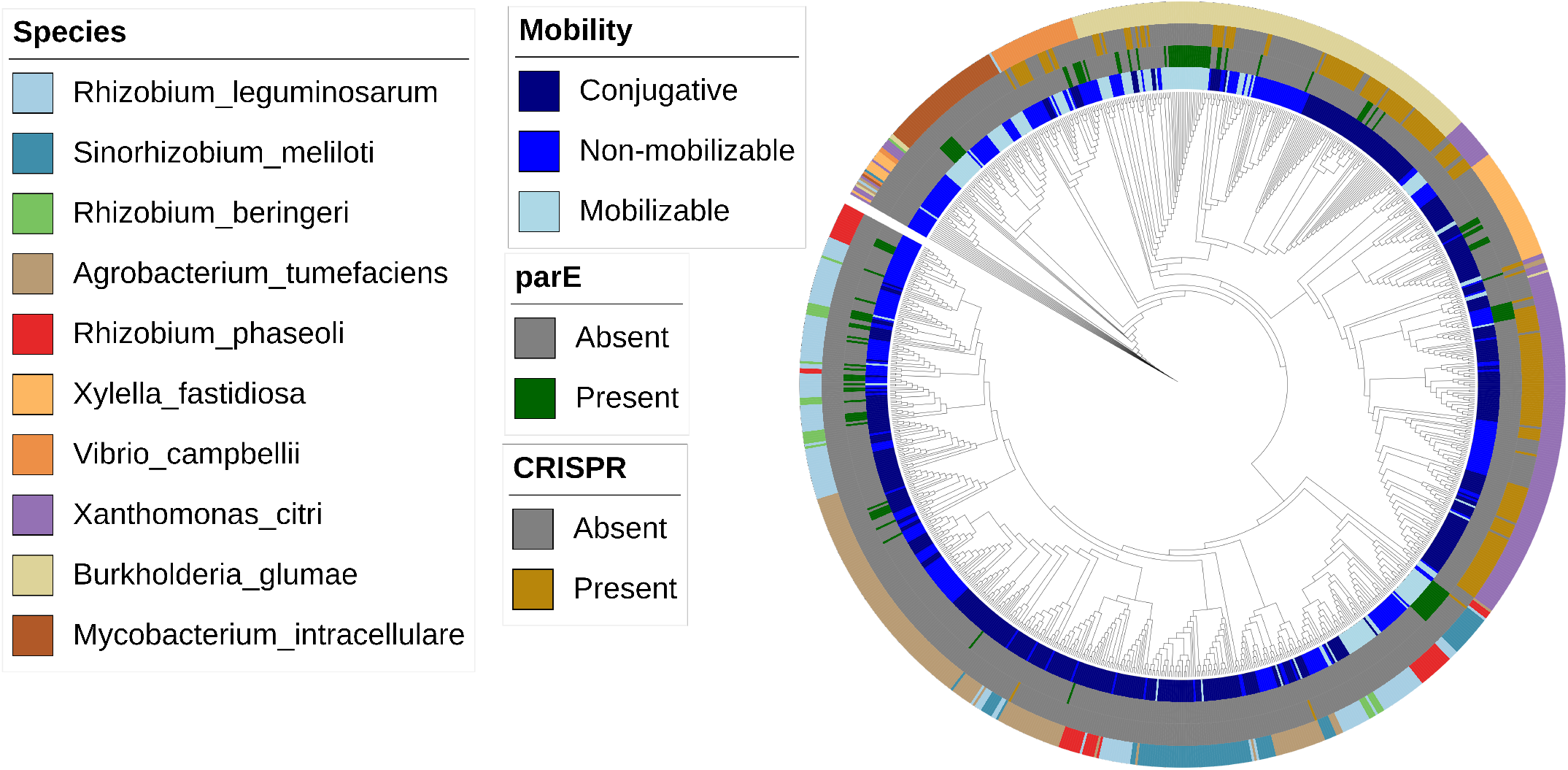
Genetic co-occurrence of *parE* and CRISPR spacers. Phylogenetic tree depicting clustering of sampled plasmid sequences for species analyzed based on gene presence-absence. Color-coded rings around the tree indicate source species, CRISPR target presence/ absence, *parE* predicted presence/ absence and predicted plasmid mobility for each plasmid included.

### 2.4 An agent-based model predicts population dynamics from biophysical parameters

Finally, we sought to synthesize our single-cell (Figs. 3 and 4) and population-level (Figs. 1 and 2) measurements into a biophysical model and assess its ability to explain the observed population dynamics. In order to incorporate spatial and single-cell variability effects, we built a CellModeller-based ([38]) agent-based model, where each cell is an “agent” whose internal state and interactions with other cells evolve according to defined rules. These rules are encoded in two modules: first, a physics module governing the spatial distribution of cells and second, a biochemistry module dealing with molecular events such as plasmid conjugation, replication and degradation (Supplementary Fig. S3). This biochemistry module is a custom implementation of the Gillespie algorithm [39] (Appendix B).

We initialized simulation runs by mirroring exactly experimental data at *t* = 0 (Figs. 6AB) and simulated according to the defined rules. Using measured conjugation rates (Fig. 3), and in the absence of a latent period for onward transmission by new transconjugants, the simulated dynamics (dashed lines) initially match the experiment for Δ*parE* RP4-0-target, but the transconjugant fraction soon exceeds the experimentally-observed fraction (Fig 6D). Altering *k*_*conj*_ could not capture observed dynamics (Supplementary Fig. S4A). Including our estimated latent period (Fig. 3C), produced excellent agreement between simulation and experiment (Fig. 6E), highlighting the importance of spatial factors for population dynamics.

**Fig. 6.**
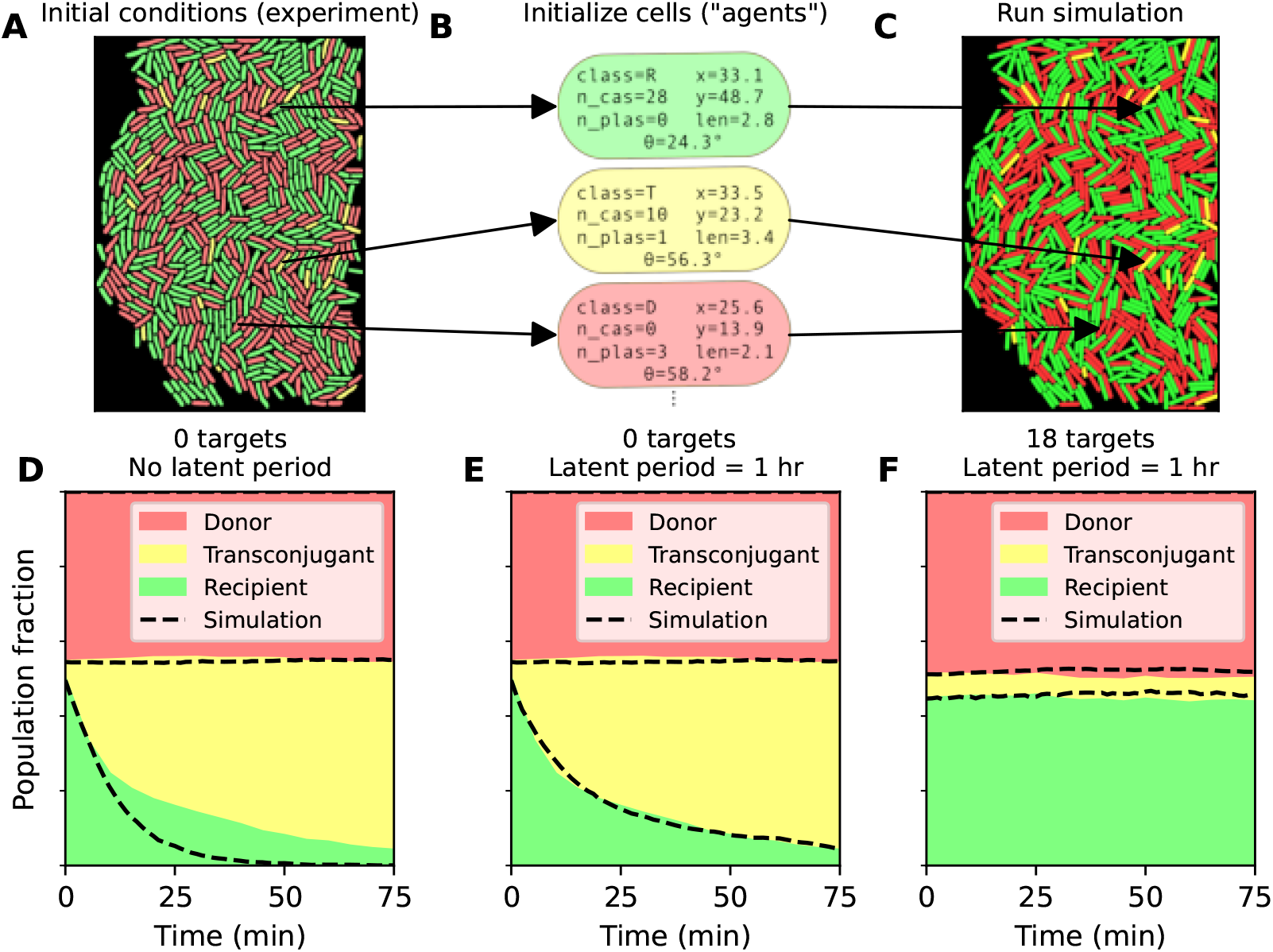
Agent-based model of plasmid population dynamics. **(A)** Initial conditions from t=0 timepoints in experiments are used as starting point for simulation runs. **(B)** For all detected cells, individual cells or “agents” are initialized with identical cell class (donor (D), recipient (R), transconjugant (T)), Cascade concentration, number of plasmids, and spatial positions as corresponding experimentally-detected cells. Three such “agents” are shown here for illustrative purposes. **C** Simulations are then run as described from initial conditions. **(D)** For 0-target plasmids, conjugation is faster in simulations (dashed lines) than in experiments in the absence of a latent period for conjugation. **(E)** The implementation of a 1 hour latent period yields excellent agreement between simulation and experiment. **F** Simulation vs. experiment for 18 target plasmid, showing good agreement when *k*_*clear*_ = 1.2 h^−1^. A complete list of parameters is found in Table S4.

Ideally, for CRISPR-targeted plasmids, another “zero parameter fit” could be achieved by simply incorporating the observed per-Cascade *k*_*clear*_ = 0.66 h^−1^ and single-cell Cascade copy numbers. Simulating the Δ*parE* 18-target case, where interference is strong, yielded a transconjugant fraction about 5 % greater than experimental observations (Supplementary Fig. S4B). To determine which *k*_*clear*_ values are consistent with experimental data, we applied Approximate Bayesian Computation (ABC) [40]. ABC performs many simulations with different parameter values, and accepts or rejects them depending on how well simulation and observed data agree (Supplementary Fig. S4CD). Here, we used the relative transconjugant fraction *T/*(*T* + *R*) to compare the two. We found that accepted simulation runs had a mean *±* standard deviation of *k*_*clear*_ = 1.5 *±* 0.3 h^−1^ per Cascade (Supplementary Fig. S4E) and good agreement with experimental results was found using *k*_*clear*_ = 1.2 h^−1^ (Fig. 6F). A complete list of parameter values is found in Supplementary Table S4. Taken together, these results show that population dynamics can successfully be predicted by spatially-explicit agent-based models based on measured biophysical parameters.

## 3 Discussion

Here, we have explored plasmid population dynamics at both the population and single-cell level, describing the role of spacer target number, Cascade expression and search time, conjugation rates, plasmid transmission latency, and toxin-antitoxin (TA) system presence. Together, these comprise a synthesis of multi-scale measurements describing the ability of CRISPR to nudge plasmid conjugation dynamics and impact overall population dynamics.

Plasmids, on average, have low spacer target numbers [41] and our bioinformatic analysis corroborates this. In addition, the wild-type level of CRISPR-RNA production in *E. coli* is very low under known growth conditions [23, 42]. Nonetheless, the wild-type Cascade constituents are capable of providing CRISPR defense when expressed [23, 42]. Similarly, we observed limited RP4 clearance when target number or expression level were low, but efficient interference when the plasmid was targeted by all spacers and Cascade was induced, holding the percentage of transconjugants at less than 10% of recipients. Acquiring more spacer targets for any single plasmid increases the clearance capability without increasing expression burden but also effectively diluting the effect of other spacers present in the genome. This increases the expression burden required to clear MGEs targeted by other spacers in the genome. This concept has been explored previously in modeling viral attack, predicting optimal spacer number [43]. Modifying spacer number may provide fine-tuning of the “nudging” capability of CRISPR interference based on ecological niche requirements. Such fine-tuning can be achieved in response to phage in days [14]. The relationship between interference and target number is also valuable knowledge for the development of simulation models and synthetic biology applications. Such applications include *e*.*g*. safeguarding against plasmid uptake and, acquisition and spread of antibiotic resistance [44].

In some circumstances, highly effective clearance proved a pyrrhic victory. RP4 encodes the *parDE* plasmid addiction mechanism which targets gyrase and ultimately decreases cell viability [45, 46]. Transconjugant cells, with established plasmids, which subsequently cleared exhibited elongated phenotypes typical of DNA damage, ultimately reducing the combined transconjugant/recipient/cleared transconjugant population. Such selective pressure against CRISPR-spacer TA co-occurrence has been noted previously [47]. Given the potential cost of expressing CRISPR defense (Fig. 4BC) [48–50], it is likely unrealistic that CRISPR defense is constitutively strongly expressed in wild populations. It is more likely that expression is initiated as required or that expression stochasticity allows populations to capitalize on a few cells with high expression. Such an assumption that most cells have low CRISPR expression most of the time makes toxin accumulation before plasmid clearance more likely, increasing the countering capabilities of TA systems. Indeed, our bioinformatic assessment suggests that CRISPR targeting is less prevalent for *parE* -containing plasmids but is clearly far from redundant in countering TA defense given its prevalence in plasmids with other toxins.

Cell-to-cell variability and its effect on population dynamics has aroused interest in recent works [21, 22, 26, 51]. The limited plasmid clearance observed here at “wildtype-like” levels (one spacer target, *≈* 100 Cascade/cell) paints a questionable picture of CRISPR efficacy against plasmids - only high target numbers could effectively clear cells. This can be countered by higher Cascade expression levels but such expression may be costly; studies have highlighted the potential costs associated with CRISPR immunity [48, 49]. Equally, the absence of immunity is a way for bacteria to obtain new genetic material [52]. In our study, we observed a linear decrease in growth rate with increasing Cascade expression levels. Many reports study CRISPR dynamics *in vitro* with relatively high resource availability [7, 21, 25, 26], which would mask potential plasmid maintenance and CRISPR interference costs. Other studies take a genetic outlook [17–20, 24] and are thus blind to environmental factors. Nonetheless, experimental [7] and theoretical [53] work has shown and explored the importance of plasmid maintenance and acquisition cost. Cascade expression stochasticity, again, may allow CRISPR to always maintain a few cells with and without plasmids, consequently upholding the ability to nudge the population towards clearance or maintenance depending on cost and benefit.

By leveraging single-cell tracking and neighbor analysis, we were able to measure key biophysical parameters: namely, Cascade search time, conjugation rate per neighboring donor cell and transconjugant latent period. Our estimate of Cascade search time is 90 minutes, in agreement with other research [25]. Conjugation rate was measured at 2.4 conjugations per hour per donor neighbor. Conjugation rate curves are well-fit by an exponential function, implying that conjugation rate is limited by a single rate-limiting step: plausibly, the conjugative pilus’ search for recipient cells [54]. We also found the latent period between plasmid acquisition and subsequent transfer to be 50 minutes; much slower than conjugation. As exemplified by our agent-based model, the estimation of such single-cell, spatially aware parameters is key for the understanding, accurate prediction and potential manipulation of population level plasmid dynamics.

Taken together, our results suggest that individual cells are the subject of a delicate trade-off between plasmid cost, plasmid benefit, defense expression, the cost of plasmid clearance, spacer acquisition and spatial considerations; neighboring cells that can influence the rate of plasmid acquisition. If CRISPR defense is expressed too little, there is limited interference. With moderate expression, and sufficient spacers, interference can be efficient. For fewer targets, expression must be higher to obtain interference, but the burden of expression may become relevant in certain environments. Furthermore, the ability of plasmids to fight back with toxin-antitoxin systems means that plasmid acquisition may be especially risky when the cost of clearance is greater than the burden of defense expression or plasmid maintenance. Maintaining a range of Cascade expression levels, coupled with varied spacer acquisition may maintain cells at all points along this continuum and subsequently be prepared for all eventualities.

## 4 Methods

### 4.1 Cloning and strain construction

#### 4.1.1 RP4 variant construction

RP4 variants were constructed by first performing the following modifications to plasmid pMM441 (a mobilizable plasmid containing an RSF1010 origin of replication and oriT, Cas9, and a guide RNA): (1) replacement of Cas9 with SCFP3A; (2) replacement of Cas9 crRNA with the pMT1 *parS* site; (3) insertion of CRISPR-Cascade targets. The part of the resulting plasmid containing a chloramphenicol resistance gene, pMT1*parS*, and Cascade targets was then PCR amplified and inserted into RP4 using recombineering.

Each of these steps will now be described in further detail. To replace Cas9 with SCFP3A, pMM441 was digested with restriction enzymes NotI and XhoI (Thermo Fisher) which flank the Cas9 coding sequence. pMM441 was a gift from Timothy Lu (Addgene plasmid #61271). At the same time, SCFP3a from plasmid p15a-Cas9deg-SOS-amp (gift from Elf lab) was PCR amplified using primers DJ001 and DJ002 which contained NotI and XhoI sites in their 5’ ends. The resulting PCR product (containing the linearized plasmid minus Cas9) was digested with NotI and XhoI, ligated into the digested pMM441 vector using T4 DNA ligase (Thermo Fisher), purified, and transformed into chemically-competent TOP10 cells, resulting in plasmid pDJ301. To replace the Cas9 crRNA with pMT1parS, pDJ301 was PCR amplified with primers DJ009 and DJ010, and a *parS* site was PCR amplified from plasmid pBlueDSBarmsparSMT1-BglII-frtCmR (gift from Elf lab) using primers DJ011 and DJ012. The resulting parS fragment was inserted into the vector using the HiFi DNA Assembly Kit (New England Biolabs), resulting in plasmid pDJ302. Next, three different derivatives were constructed from pDJ302, containing 0, 1, or 18 targets for Cascade. To create plasmids pDJ310 and pDJ311 (containing 0 and 1 target, respectively), pDJ302 was PCR amplified with primer pairs DJ003/DJ004 and DJ005/DJ006, respectively. Primers DJ003/DJ004 contained 5’ “floppy ends” such that, when the resulting linear product was circularized by ligation, a CTT PAM site adjacent to either a 32 bp randomized DNA sequence would be inserted into the plasmid. Similarly, primers DJ005/006 were designed to insert a protospacer corresponding to spacer 1 from the E. coli CRISPR1 array.

At the ParS/ParB-mCherry system exhibited good functionality in identifying plasmid-carrying cells, using CFP expression to identify plasmid-carrying cells was not necessary, and CFP could be deleted. To delete SCFP3a from pDJ310/311, the plasmids were PCR amplified with primers DJ075/DJ076, which were designed not to amplify the region containing SCFP3a and its promoter. The resulting linear products were phosphorylated using T4 PNK, blunt end ligated with T4 DNA Ligase, cleaned up, and transformed into TOP10 chemically competent cells (Thermo Fisher), thus creating plasmids pDJ320 (no targets) and pDJ321 (1 target). To create plasmid pDJ323 (containing all 18 targets from the *E. coli* CRISPR arrays 1 and 2), pDJ320 was PCR amplified and linearized using primers DJ019/DJ115 to create a vector backbone. At the same time, plasmid pTarget (gift from Brouns lab), containing all 18 targets from *E. coli* CRISPR arrays 1 and 2, was amplified using primers DJ022/116. The primers were designed with overlaps appropriate to HiFi DNA Assembly. The two fragments were assembled using HiFI DNA Assembly, purified, and transformed into TOP10 chemically competent cells (Thermo Fisher Scientific).

To create the *wt parE* RP4 variants(also referred to as +*parE*) used in this work, a cassette containing (1) chloramphenicol resistance (2) pMT1*parS* (3) 0, 1, or 18 CRISPR targets was amplified from plasmids pDJ320, pDJ321, and pDJ323, respectively, using primers DJ066/DJ067 with AccuPrime Pfx DNA Polymerase (Thermo Fisher). These primers contained homology arms appropriate to lambda red recombination into RP4, upon which the cassette replaced the divergently transcribed tetA-tetR genes in RP4. The cassettes were integrated into RP4 using lambda red recombination and restreaked three times to ensure segregation of recombineered RP4 variants. Clones were streaked on tetracycline plates to ensure complete segregation (since segregated clones are sensitive to tetracycline). The resulting strains were EL4120, EL4121, and EL4122.

To create the Δ*parE* RP4 variants used in this work, we chose to back up a few steps and start over from *wt* RP4. Our approach was first to delete *parE* from *wt* RP4 using the DIREX method [55], and then re-introduce the cassette containing CRISPR targets and the *parS* site. To delete parE, we amplified the *AcatsacA* cassette from genomic DNA isolated from strain DA42825 (gift of Andersson lab) with primers DJ128/DJ062 and DJ063/DJ129 which contain appropriate overhangs to replace *parE* with the *AcatsacA* cassette while creating 40 bp direct repeats immediately outside of the cassette. The *AcatsacA* cassette contains the *cat* gene for chloramphenicol resistance as well as the *sacB* gene imparting sucrose sensitivity. The cassette also contains two copies of the *amilCP* blue chromoprotein as inverted repeats at either end of the cassette. It is necessary to amplify the cassette in two separate reactions due to the inverted *amilCP* repeats; the two fragments contain a 277 bp overlap in the *cat* gene which is subsequently recombined *in vivo* under lambda red.

The amplified cassette was eletroporated into strain EL3316 which contains *wt* RP4 as well the pSIM5-tet lambda RED recombinase plasmid. The resulting unstable intermediate RP4-parE::AcatsacA has parE replaced by the AcatsacA cassette. The cassette can be maintained by propagation in chloramphenicol-containing media. To resolve the unstable intermediate and create the Δ*parE* RP4, cells were streaked on sucrose-containing plates to select for cells in which recombination occurred. The designed direct repeats as well as the inverted *amilCP* repeats recombine to yield a scarless deletion of *parE* (see ref. [55] for a pedagogical explanation of this technique). Primers were designed to perform an in-frame deletion with the first 10 and last 10 amino acids of ParE left intact.

With the RP4-ΔparE plasmid in hand, the cassettes from plasmids pDJ320, pDJ321, and pDJ323, were amplified as before and integrated into RP4-ΔparE using lambda RED recombineering (also as before), yielding strains EL4519-4522 with 0, 1, and 18 CRISPR targets, respectively. The three RP4 versions were then conjugated into strain EL4009 to create strains EL4533-4535 used as donors in the paper.

DNA cleanup steps were performed with the Monarch PCR and DNA Cleanup Kit (New England Biolabs) unless otherwise specified. PCR steps were performed with Q5 DNA Polymerase unless otherwise specified (New England Biolabs).

#### 4.1.2 Donor strain construction

As *E. coli* will opportunistically use arabinose as a carbon source depending on the presence of alternative carbon sources, we elected to delete the *araBAD* operon in experimental strains to ensure that arabinose would function only as an inducer and not as a carbon source. The *araBAD* operon was deleted and replaced with a kanamycin resistance marker in strain MG1655 using lambda red recombination, yielding strain EL4008. The araBAD deletion was subsequently transferred to strain EL904 (MG1655 rph+) using P1 transduction, yielding strain EL4009. Finally, the three *wt parE* RP4 variants containing 0, 1 and 18 targets were respectively conjugated from strains EL4067, EL4068, and EL4069, into strain EL4009, yielding strains EL4020, EL4021, and EL4022. The Δ*parE* variants were similarly conjugated into strain EL4009 yielding strains EL4533, EL4534, and EL4535.

#### 4.1.3 Recipient strain construction

Creation of the recipient strain EL4012 from strain MG1655 involved a number of steps: (1) creating an N-terminal fusion of SYFP2 with Cas8e; (2) P1 phage transduction of mCherry-ParB into the *gtrA* locus; (3) replacement of the Pcas3 promoter with the Placuv5 promoter; (4) replacement of the Pcascade promoter with the araBp8 promoter (5) deletion of araBAD operon. In replacing the *cas3* and Cascade promoters, we are following the appproach of Datsenko *et al* [56]. For recipient strain El4014, we also deleted the CRISPR1 and CRISPR2 arrays.

##### Creation of N-terminal SYFP2-Cas8e fusion

To create the fusion, we used the Dup-In method to which enables creation of markerless, scarless fusions [55, 57]. This technique involves first creating a 500 bp duplication of *sfyp2* flanking a selectable/counter-selectable *Acatsac1* cassette, which contains the *cat* gene for chloremphenicol resistance and the *sacB* gene from *Bacillus subtilus* conferring sucrose sensitivity. This construct has previously been created in the Dan Andersson lab as strain DA43403. We amplified the *syfp2* duplication-*Acatsac1* cassette from genomic DNA isolated from strain DA43403 (gift of Andersson lab) using primers with homology such that the product would recombine in the N-terminal position relative to *cas8e*. Due to the duplicated *syfp2* segments, the entire construct cannot be amplified at once. Rather, two separate PCRs were performed using primers DJ062/DJ065 and DJ063/DJ064 respectively. Primer DJ065, containing homology to the N-terminus of *cas8e*, was designed such that *sfyp2* would lack a stop codon, with an 18bp linker sequence (as in Vink *et al* [25]), and deleting the start codon of *cas8e*. The two PCR products overlap by 277 bp in the *cat* gene such that they will recombine to form the correct, complete construct using lambda red recombination (see Figure 4 in [57]).

The result of this lambda red recombination was strain EL3177. As long as this strain is propagated in chloramphenicol-containing media, the unstable duplication will be maintained. The construct was then transferred to MG1655 *rph+* using P1 phage transduction. After isolation of a successful transductant, the strain was grown on sucrose-containing plates to select for colonies in which the flanking *syfp2* duplications have recombined, eliminating the *Acatsac1* cassette, and yielding an N-terminal fusion as desired (strain EL3178).

##### P1 phage transduction of mCherry-ParB

A cassette containing an mCherry-ParB fusion protein expressed from a medium-strength constitutive promoter, as well as a spectinomycin resistance marker, was transduced from strain EL851 (gift of Elf lab) to strain EL3178 to create strain EL3202. The cassette is integrated in the *gtrA* locus.

##### Replacement of Pcas3 promoter with Placuv5 promoter

The Pcas3 promoter was first replaced with the *Acatsac1* cassette (described above). To accomplish this, the *Acatsac1* cassette was PCR amplified with primers DJ079/DJ080 which contain appropriate homology flanking Pcas3. This product was integrated into the chromosome using lambda red recombineering, thus replacing Pcas3 with the *Acatsac1* cassette, and yielding strain EL3654. The PlacUV5 promoter was ordered as a gBlock (Integrated DNA Technologies) also containing homology flanking the now- deleted Pcas3 promoter. This gBlock was then integrated into the genome again using lambda red. Successful clones were identified by sucrose counter-selection (as correct clones will delete the *Acatsac1* cassette), yielding strain EL3658.

##### Replacement of Pcascade promoter with araBp8 promoter

A similar two-step approach was employed to replace the Pcascade promoter. First, the Pcascade promoter was replaced by the *Acatsac1* cassette using lambda red recombination, yielding strain EL3661. Second, the araBp8 promoter was PCR amplified from strain KD263 (gift from Severinov lab) using primers DJ092/DJ093 which also contain homology flanking the deleted Pcascade promoter. This PCR product was integrated into strain EL3661 using lambda red recombination and correct clones were isolated using sucrose counter-selection, yielding strain EL3701.

At this point we had achieved the desired construct, and several rounds of lambda red recombination had been performed. To mitigate the risk of off-target mutations accumulating elsewhere in the genome, we performed a final step of transducing the modifications in the vicinity of the Cascade operon into a clean background strain. To create a clean background strain, we transduced the *Pcas::Acatsac1* construct from strain EL3654 into strain EL3202 using P1 phage transduction, yielding strain EL3946. This strain has not had any lambda red recombinations performed in it, only P1 phage transductions. Finally, the modifications from strain EL3701 were transduced into strain EL3946 using P1 transduction, which was possible since clones with the correct transduction could be idenfitied by sucrose counter-selection (only successful transductants can grow on sucrose). This yielded strain EL3957.

##### Deletion of *araBAD* operon

Again to ensure that arabinose would be used only as an inducer and not a substrate for growth, the *araBAD::kanR* deletion was transduced from strain EL4008 into strain EL3957, yielding strain EL4012 used as recipient in the manuscript.

##### Deletion of CRISPR1 and CRISPR2 arrays

For deletion of the CRISPR arrays, we again used the DIREX method to create markerless, scarless deletions [55]. For each array, we kept the leader and terminator sequences intact while deleting all spacers from the array. To delete the CRISPR1 array, we amplified the *AcatsacA* cassette from genomic DNA isolated from strain DA42825 (gift of Andersson lab) with primers DJ098/DJ062 and DJ063/DJ099 which contain appropriate overhangs to replace CRISPR1 with the *AcatsacA* cassette while creating 40 bp direct repeats immediately outside of the cassette, and also creating a 277 bp overlap in the *cat* gene for recombination of the two fragments as described above. The products were integrated into strain EL3701 using lambda red recombination, and after confirmation of the correct clone, the unstable intermediate was resolved by growth on sucrose-containing media yielding the desired deletion of CRISPR1. An identical procedure was followed for CRISPR2, with the *AcatsacA* cassette being amplified by primer pairs DJ100/DJ062 and DJ063/DJ101, yielding strain EL3898 with both arrays deleted.

To avoid accumulation of off-target mutations due to multiple lambda red rounds, we transduced this construct into strain EL3946 using P1 phage transduction. Since the CRISPR1 and CRISPR2 arrays are separated by around 20 kb, and P1 transduction transfers approximately 100kb at a time, it is possible to transduce both mutations at once, subject to sequence verification. Finally, the *araBAD::kanR* deletion was transduced from strain EL4008 to create strain EL4014, the desired final construct.

### 4.2 Microfluidic assays

Strains were grown overnight in LB broth (tryptone 10 g/L, yeast extract 5 g/L, NaCl 10 g/L) from single colony with appropriate antibiotic addition (spectinomycin 15 *µ*g/L for recipient strains or chloramphenicol 12.5 *µ*g/L for donor strains) shaking 200 rpm 37^*°*^C. Overnight cultures were diluted 20 *µ*L into 10 mL M9 media in 50 mL falcon tubes. Final M9 media composition is as follows (Na_2_HPO_4_ 12.8 g/L, KH_2_PO_4_ 3 g/L, NaCl 0.5 g/L, NH_4_Cl 1 g/L, CaCl_2_ 100 *µ*M, MgSO_4_ 2 mM, x1 RPMI 1640 amino acid solution 1640 (Sigma R7513), glucose 0.4%, pluronic F-127 85 mg/L (Sigma-Aldrich P2443)), where inducers were included in the media these are final concentration (IPTG (isopropyl *β*-D-1-thiogalactopyranoside) 1 mM, arabinose 0.2%). Diluted cells were returned to shaking incubation for three hours; when cells were in exponential growth cells. At this point cells were mixed 1:1 (donor:recipient) by volume and loaded into microfluidic chips for imaging. The device used has two parallel row of traps separated by a central barrier. This allows loading of two different strain combinations in each experiment. For most experiments +parE and -parE variants of the same donor strain were run concurrently. For experiments with delayed induction, the +/- CRISPR array-containing recipient strains were run concurrently with one donor strain.

The microfluidic device used here is a modified version of the device used in Baltekin et al 2017[58] with a 1000 *µ*m channel height and large rectangular cell traps measuring 48 *µ*m x 60 *µ*m. Cells were loaded into these traps immediately after mixing under 0.4-0.6 bar pressure. After cells were loaded fresh M9 media was flown into the device at 0.2 bar and continued for the duration of each experiment. Imaging setup was composed of a Nikon Ti2 equipped with Teledyne Kinetix camera, 100x Ph3 objective (CFI Plan Apo lambda 100X/1.45 oil. MRD01905), 1.5x internal magnification selected (for a total magnification of 150x), Lumencore SPECTRA light engine and filters configured for mCherry (No excitation filter, Chroma ET645/75M emission filter and Chroma ZT594rdc dichroic mirror) and SYFP2 (Chroma Z514/10 excitation filter, Semrock FF01-542/27M emission filter and Semrock BrightLine Di02/R514 dichroic mirror) fluorescence imaging. The microscope stage was encased in Okolab H201-ENCLOSURE hood and H201-T-UNIT-BL temperature control unit to maintain a constant 30^*°*^C throughout the experiments. Cells were imaged in phase contrast (100 ms exposure), mCherry fluorescence (200 ms), SYFP2 fluorescence (100 ms) channels over a period of 6 hours. Phase, mCherry and SYFP2 images were taken at 1.25, 5, and 30 min intervals, respectively.

### 4.3 Image analysis

The image analysis performed in this work is based on the following foundational steps: segmentation, cell tracking, dot (i.e., plasmid) detection, and cell classification. These steps are followed by downstream analyses such as plasmid clearance rates, cell death rates, growth rate computations, etc. Foundational steps as well as downstream analyses were integrated into a custom image analysis pipeline based on the pipeline described in ref. [59]. All code used to perform the analyses is available upon request.

Segmentation was performed on phase contrast images using the Omnipose software package [60], which uses a U-net-based neural net architecture to perform morphology-independent cell segmentation. Custom neural nets were trained for this project using manually-curated segmentation of cells grown in microfluidic devices as ground truth. Cell tracking was performed using the Baxter algorithm [35]. Dot detection was performed using the radial symmetry algorithm; detected dots were subsequently refined by fitting a Gaussian to each detected dot.

To classify cells as donors, recipients, and transconjugants, a two-step process was used. Recipients and transconjugants express ParB-mCherry, whereas donors have no signal in the mCherry channel. Cells were first classified as “mCherry positive” or “mCherry negative” by thresholding, where the threshold was chosen to minimize the standard deviation of fluorescence signal intensity within the respective “mCherry positive” and “mCherry negative” groups. “mCherry negative” cells can then immediately be identified as donors. “mCherry positive” cells containing detected mCherry dots (corresponding to ParB-mCherry-tagged RP4 plasmids) are classified as transconjugants, whereas “mCherry positive” cells not containing detected mCherry dots are classified as recipients. For analyses involving cell tracking across multiple frames, transconjugant cells must have two consecutive frames without detected dots in order to revert to recipient status. This is to avoid false clearance events resulting from failure to detect plasmids in a single frame (which can plausibly occur due to RP4’ s relatively low copy number and the fact that fluorescence imaging takes place in a single z-axis plane, so that plasmids may be undetected due to being out of focus).

Cell neighbors were determined using morphological operations on segmentation masks. For each segmented cell, the segmentation mask was cropped to the bounding box of the cell plus 15 pixels in each dimension. (Pixel size is 6.5 *µ*m*/*150 *≈* 0.043 *µ*m per pixel). The cell of interest was dilated by six pixels, and the other cells within the cropped region were subsequently dilated by six pixels, one at a time. When the central cell’ s dilated mask overlapped by 40 pixels or more with another cell’ s dilated mask, the cells were determined to be neighbors. We experimented with different thresholds and found empirically that 40 pixels was a reasonable threshold to match our judgements by eye of which cells were neighbors.

#### Latent period lineage selection

For conjugation latent period calculations, we selected recipient cell lineages based on the following criteria: 1. no donor neighbors prior to conjugation 2. no transconjugant neighbors for at least the first 10 minutes of the lineage (to ensure that conjugation is performed by “new” transconjugants undergoing the latent period) 3. starting from the first frame with a transconjugant neighbor, a transconjugant neighbor must be present in at least 80% of frames until conjugation.

### 4.4 Cascade copy number calculations

*E. coli* strain EL4269 was used as a basis for the calculation of Cascade copy number. This strain contains SYFP2 fused to lacI along with a single genomic lacI-binding site Osym. The strain was induced and prepared in the same way as cells used in experiments. After growth to exponential growth phase cells were mounted on agarose pads (1.5% agarose, M9 + inducers as described above). Cells were imaged on the same microscope set-up as described above but with varying exposure times and light intensity settings (1000 ms, 10% intensity and 500 ms 40% intensity). Circular ROIs were drawn around visible dots within cells (representative of a single bound Cascade molecule) and a background measurement taken inside the same cell away from the dot position with the same size and shape ROI. By averaging background subtracted measurements over biological and technical repetitions we obtained a value of 6382 and 2222 fluorescence units per Cascade molecule under (1000 ms and 500 ms) imaging conditions, respectively. Next we used constitutive SYFP2 expressor, strain EL2788 imaged under the same conditions as the LacI fusion strain. Manual cell counting, automated masking with Fiji auto-threshold function and intensity measurements allowed us to calculate the copy number as approximately 23 Cascade copies per cell across all replicates. We than imaged the constitutive expressor in microfluidic traps under experimental imaging conditions. Given the now known copy number per cell we were able to calculate 526 total arbitrary fluorescence units per Cascade-SYFP2 molecule under experimental imaging conditions. This value was then used to calculate copy number from background subtracted fluorescence values in experiments.

### 4.5 Bioinformatics analysis

CRISPR spacer sequences (http://crispr.genome.ulaval.ca/) and *parE* sequences (XCD26522.1) were used to search plasmid sequences[36] using local blastn and tblastn respectively. For spacer sequence hits those with 100% sequence identity over 90% of the sequence length were retained. For *parE* sequences, hits with evalue *<* 1 were retained and considered *parE* -like sequences for our analysis.

We constrained our analysis to plasmid host species in which there was evidence of *parE* presence. We calculated the predicted proportion of plasmids having a *parE* hit for each species and selected the top 10. The predicted proportion of plasmids was estimated using an empirical Bayes approach by applying a beta prior to the proportion for species *i* where *β* and *α* are *parE* -containing and *parE* -free plasmids across the global dataset and *n* and *x* are the total and *parE* -containing, respectively, number of plasmids.

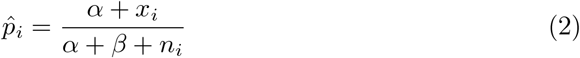

To account for plasmid phylogenetic relatedness we used a gene clustering and co-occurrence method. We first took all the amino acid sequences for annotated genes in selected plasmids and applied pairwise sequence alignment and clustering using MMseq2[61], 70% sequence identity over 80% of sequence length, to create clusters of likely gene orthologs. After this step, plasmid collections were randomly sub-sampled to a maximum depth of 100 plasmids for each species-mobility combination; giving a maximum of 300 plasmids per species. Cluster assignments were then used to calculate pairwise Jaccard distances between all plasmids selected and construct phylogenetic relationships, similar to the method described in [62]. Phylogenetic distances derived from the constructed tree were used as probability weightings to perform phylogenetic permutations (as described previously[63]) to test the significance of the relationship between toxin presence and CRISPR target presence. The methods described above were repeated for 8 additional known toxin proteins: *relE, vapC, pemK, mazF, txe, doc, CcdB, higB*.

## Supplementary figures and tables

Further supplementary information can be found in the Appendices in this document.

## Acknowledgements

We are grateful to the Brouns lab for the gift of the pTarget plasmid, the Semenova and Severinov labs for the gift of strain KD267, the Andersson lab for the gift of strains DA43403 and DA42825, the Elf lab for the gift of strain EL851, and the Lu lab for the gift of plasmid pMM441. Useful discussions and feedback were provided by Anna Knöppel, Jimmy Larsson, Johan Elf, Brian Rydgren, Filippa Nilsson, and Nathalie Balaban. DJ acknowledges support from the Swedish Research Council (grant number 2020-05137). LR acknowledges support from the Carl Tryggers Foundation (grant number CTS 21:1334). Computation and data management were enabled by resources provided by the Swedish National Infrastructure for Computing at UPPMAX.

## Author Contributions

LR performed experiments, analyzed data, and wrote analysis code. DL performed experiments, and designed and constructed strains. JC developed experimental protocols. JW wrote analysis code. AT developed agent-based simulation code. DJ analyzed data, wrote analysis code, and conceived and supervised the project.

## Supplementary Information

**Fig. S1.**
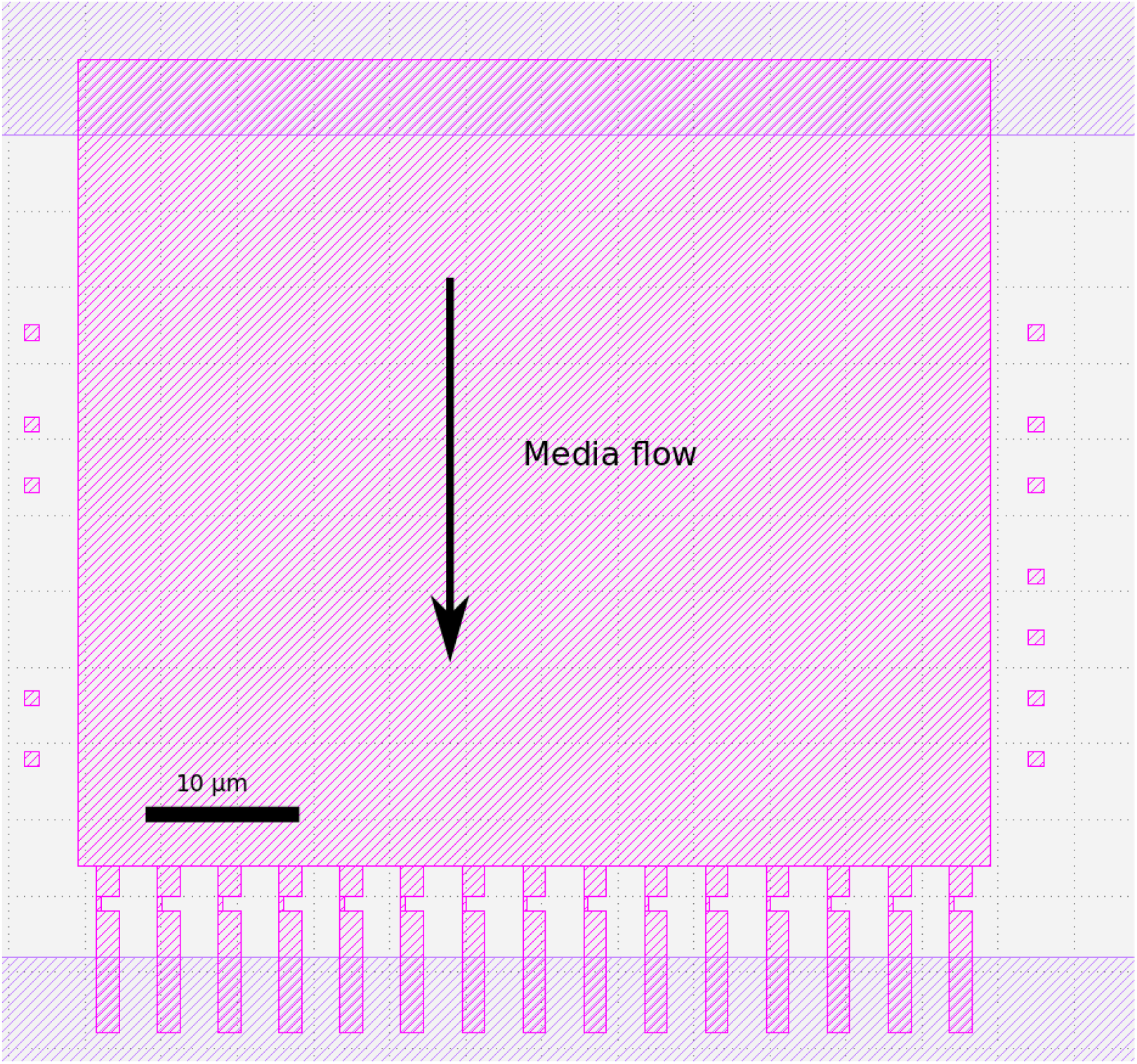
Schematic of of a single microfluidic trap/chamber. Cells occupy the rectangular area which measures 48×60 *µ*m. The channels at the bottom are 1.5 *µ*m wide with 0.25 *µ*m constrictions, enabling media to flow over the cells as indicated while preventing cells from passing through. Each chip contains 200 such traps.

**Fig. S2.**
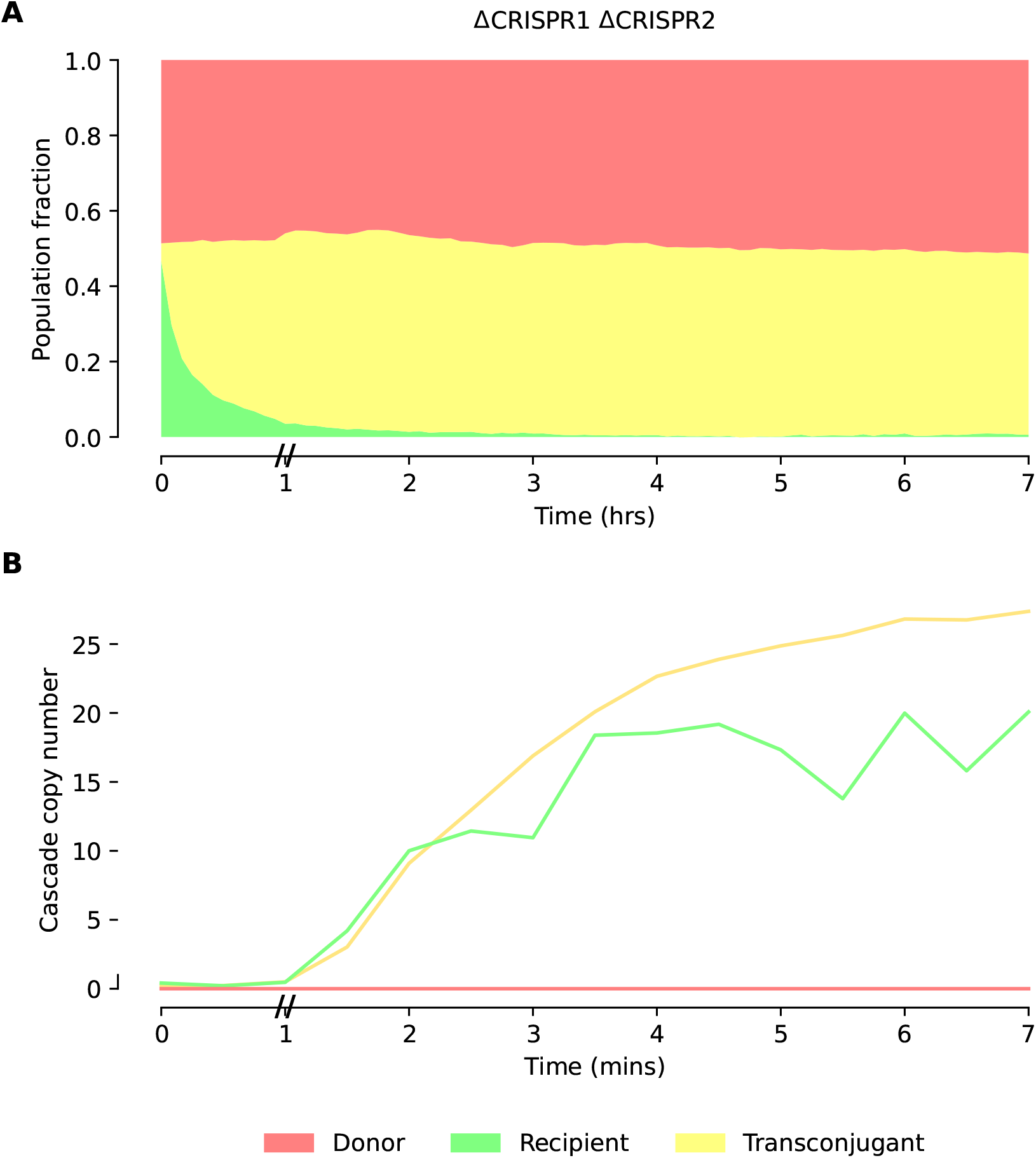
A) Mean fraction of the cell population classified as donor, recipient or transconjugant across the course of microfluidic experiments with delayed Cascade induction with 18-target RP4 plasmids without *parE*, for recipient *E. coli* strains with genomic CRISPR arrays deleted. B) Median Cascade copy number per cell across the course of the experiment. Inducers were added to the media at 60 minutes, indicated with // on axis.

**Fig. S3.**
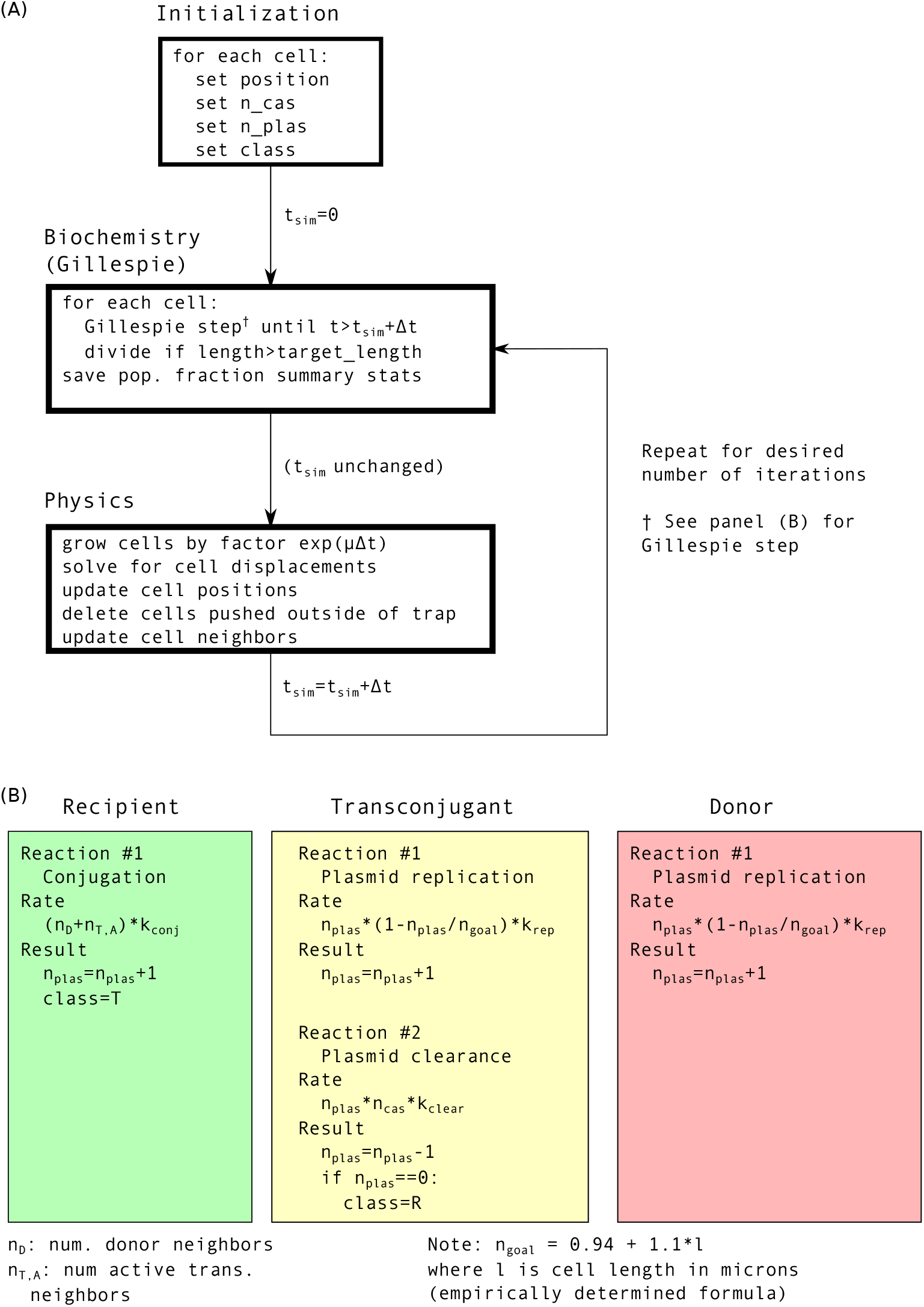
**(A)** Overview of agent-based simulation. After initialization, biochemistry and physics modules are run for each timestep Δ*t*. This is repeated for the desired number of iterations. We chose Δ*t* = 1.25 min and 50 iterations, for a total simulation length of 75 min. **(B)** Possible reactions, rates thereof, and resulting change in chemical species for each cell class in Gillespie simulation. While the physics module takes fixed timesteps Δ*t*, Gillespie simulations have variable timesteps; therefore, during each iteration, Gillespie simulations are run starting from *t*_*sim*_ until the next reaction timepoint exceeds *t*_*sim*_ + Δ*t* where *t*_*sim*_ is the current simulation time.

**Fig. S4.**
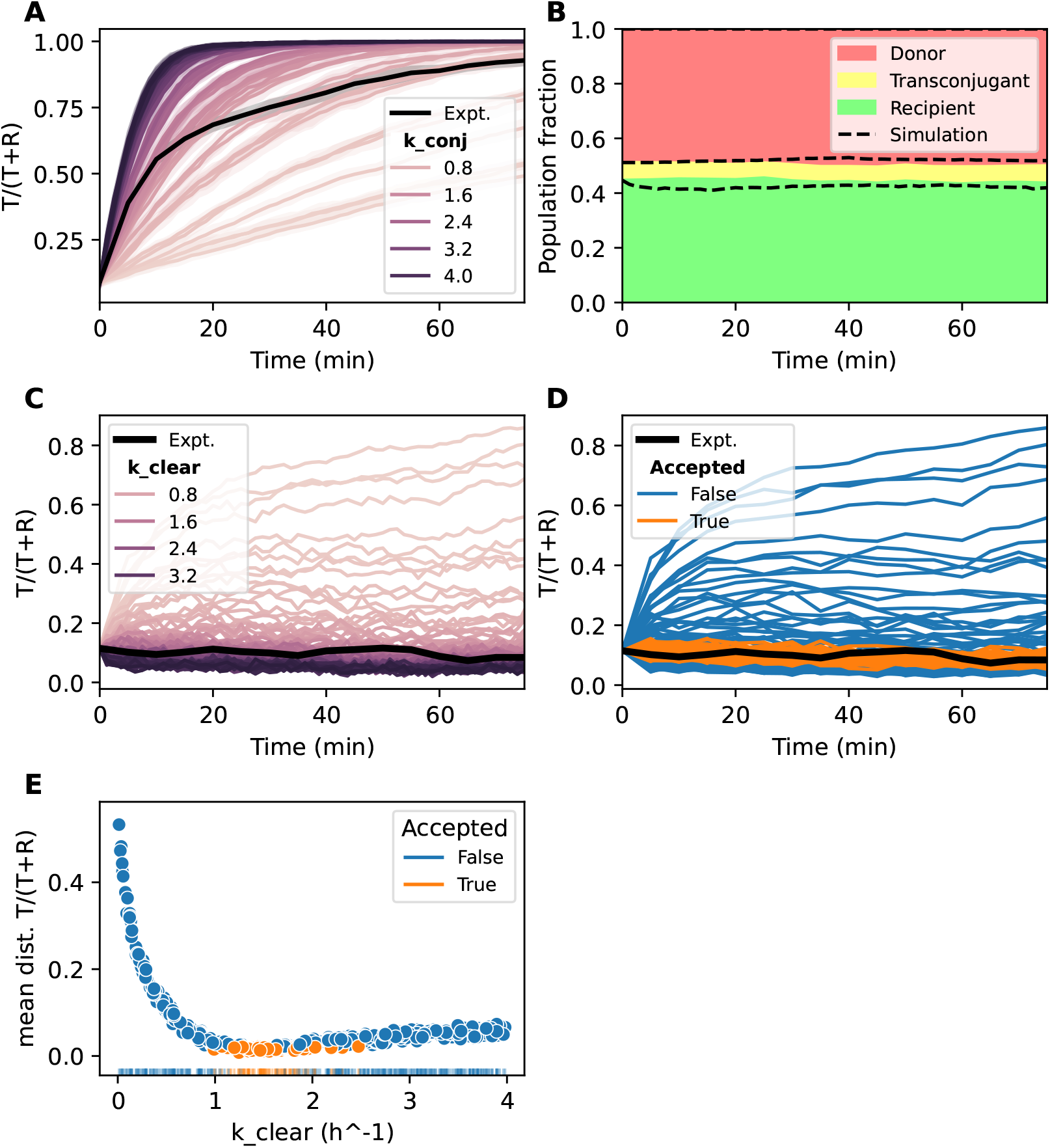
Optimizing parameter values using Approximate Bayesian Computation (ABC). **(A)** Relative transconjugant fraction *T/*(*T* + *R*) from simulations performed for 0-target case with no latent period for different values of conjugation rate *k*_*conj*_. In the absence of a latent period, no *k*_*conj*_ values match experiment at both early and late times. All rates in all panels in units of h^−1^. **(B)** Comparison between simulation (dashed lines) and experiment (solid colors) for 18-target plasmid with 1 hour latent period and *k*_*clear*_ = 0.66 h^−1^ as determined in Figure 4. The simulated transconjugant fraction is about two times the observed value. **(C)** Dependence of relative transconjugant fraction *T/*(*T* + *R*) on clearance rate *k*_*clear*_ for randomly chosen values between 0 and 4 h^−1^. **D**. ABC analysis of trajectories shown in panel C. *k*_*clear*_ values whose *T/*(*T* + *R*) curves deviate by 0.03 or less from experimental data are accepted; other values are rejected. **E**. Another way of looking at panels C and D. The mean distance in relative transconjugant fraction is shown as a function of *k*_*clear*_; acceptance status indicated by color. A “rug plot” along the x axis shows individual accepted and rejected values.

**Table S1.** Inferred plasmid clearance rates *k*_clear_, computed both naively and accounting for conjugation, false negatives, etc (see Appendix A). Two consecutive plasmid-free frames were required to call a clearance event. Rates computed over all cells (average copy number 42 Cascade per cell). To obtain approximate rate per Cascade, divide by 42. To obtain corresponding rates for “wild-type” expression levels (*≈* 125 Cascade per cell), multiply rates by 3. Corrected rates are higher because the non-negligible probability of an additional conjugation event during the two required plasmid-free frames means that the apparent clearance rate will be lower than the true clearance rate (Appendix A).

| Plasmid | Naive $k_{\text{clear}}$ ($\text{h}^{-1}$) | Corrected $k_{\text{clear}}$ ($\text{h}^{-1}$) |
| --- | --- | --- |
| RP4-0-target | 0.84 | 0.0 |
| RP4-1-target | 0.90 | 0.18 |
| RP4-18-target | 10.56 | 18.36 |
| RP4-0-target$\Delta parE$ | 1.32 | 0.9 |
| RP4-1-target$\Delta parE$ | 0.96 | 0.24 |
| RP4-18-target$\Delta parE$ | 9.36 | 16.14 |

**Table S2.**
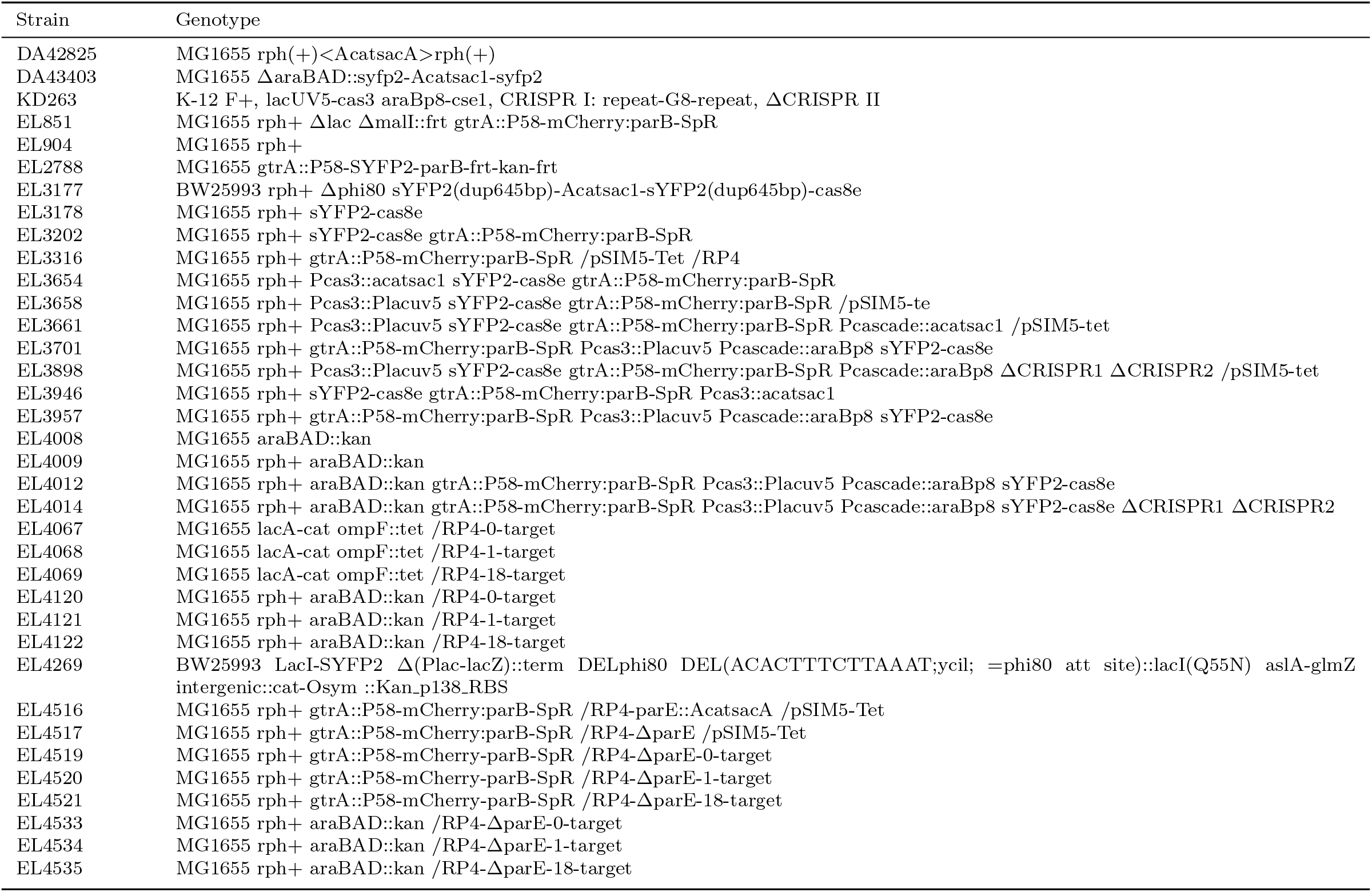
List of strains used in this work.

**Table S3.**
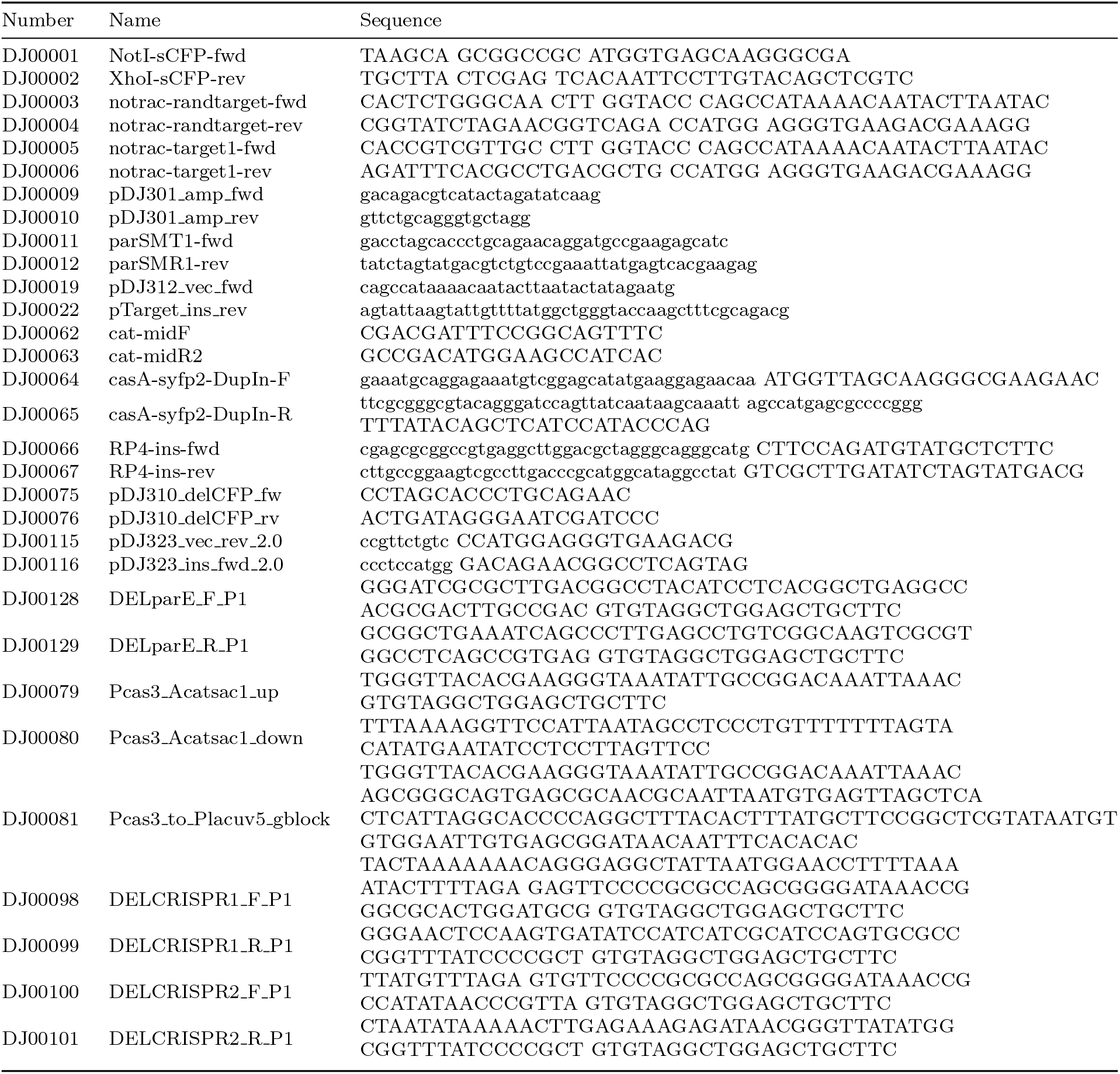
List of primers used in this work.

**Table S4.** List of parameters used in agent-based model. All rates and times in hours unless otherwise indicated.

| Parameter name | Description | Value |
| --- | --- | --- |
| kd_plasmid_division | Plasmid replication rate | 4.7 |
| max_plasmids | Maximum allowed plasmid number | 10 |
| plas_vs_length_intercept | Num plas. vs length intercept | 0.412 |
| plas_vs_length_slope | Num plas. vs length slope | 0.877 |
| plas_vs_length_sd_intercept | Num plas. vs length stddev intercept | 0.525 |
| plas_vs_length_sd_slope | Num plas. vs length stddev slope | 0.263 |
| latent_period | Latent period for conjugation | 1 |
| k1 | Plasmid clearance rate (called k_clear in text) | 0.66 |
| target_sites | Num. target sites on plasmid | 0 (Fig. 6DE) / 12 (Fig. 6F) |
| cascade_spacers | Number of spacers in CRISPR1 array | 12 |
| growth_rate | Growth rate | 1.05 |
| target_length | Target length for division | 5 |
| cell_radius | Cell radius (um) | 0.4 |
| dt | Simulation step time interval | 0.0208333333333333 |
| conjugation_rate | Called k_conj in text | 2 |

**Table S5.** Outputs relating to phylogeny-weighted permutation tests per toxin gene analyzed. For each toxin gene we present, for the host species selected, the count of plasmids with CRISPR targets and toxin-like genes as well as the plasmids with both features co-ocurring. We then present the number of times the co-occurring count is greater than or less than the observed data for 10000 permutations with associated p values and adjusted p values (Benjamini Hochberg correction)

| Gene | Accession | CRISPR_count | Gene_count | Both_count | Both_sim_greater |
| --- | --- | --- | --- | --- | --- |
| **ccdB** | AAA96062.1 | 1746 | 319 | 311 | 0 |
| **txe** | ADV60046.1 | 364 | 216 | 78 | 4451 |
| **relE** | ATP89023.1 | 1640 | 299 | 242 | 0 |
| **vapC** | AVR61228.1 | 1398 | 361 | 287 | 0 |
| **pemK** | AVV60552.1 | 1314 | 396 | 257 | 8572 |
| **higB** | AWD72426.1 | 1116 | 352 | 244 | 1823 |
| **mazF** | CAD6005292.1 | 953 | 380 | 210 | 9735 |
| **doc** | QHW11153.1 | 965 | 233 | 153 | 1534 |
| **parE** | XCD26522.1 | 223 | 121 | 15 | 9993 |
| Gene | Both_sim_lesser | p ≤ Both_count | p ≥ Both_count | p-adj ≤ Both_count | p-adj ≥ Both_count |
| **ccdB** | 10000 | 1 | <0.0001 | 1 | <0.0006 |
| **txe** | 4924 | 0.5549 | 0.5076 | 0.99882 | 0.99882 |
| **relE** | 10000 | 1 | <0.0001 | 1 | <0.0006 |
| **vapC** | 10000 | 1 | <0.0001 | 1 | <0.0006 |
| **pemK** | 1206 | 0.1428 | 0.8794 | 0.4284 | 1 |
| **higB** | 7808 | 0.8177 | 0.2192 | 1 | 0.4932 |
| **mazF** | 199 | 0.0265 | 0.9801 | 0.0954 | 1 |
| **doc** | 8075 | 0.8466 | 0.1925 | 1 | 0.4932 |
| **parE** | 4 | 0.0007 | 0.9996 | 0.00315 | 1 |

## Appendix A Estimation of plasmid degradation / CRISPR interference rates

### A.1 Purpose

We want to estimate clearance/plasmid degradation rates from time-lapse fluorescence microscopy observations of plasmid copy number. The fact that our observations take place at discrete time intervals of 5 minutes requires some consideration, as well as the possibility of additional conjugation events taking place during each 5 minute interval.

Thoughout, we will assume that we are starting with an observation of one plasmid at *t* = 0. We will also assume that 5 minutes is a sufficiently short time interval that plasmid replication cannot occur within this interval. All times are in minutes, and all rates are in units of min^−1^.

### A.2 Naive approach

If there is one plasmid at *t* = 0, and plasmids are degraded at rate *k*_*d*_, the probability of zero plasmids at *t* = 5 is given by the cumulative distribution function (cdf) of the exponential distribution with rate *k*_*d*_:

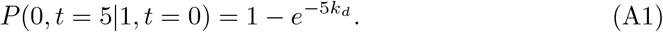

This can easily be solved for *k*_*d*_:

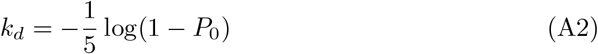

where we defined *P*_0_ = *P* (0, *t* = 5 |1, *t* = 0) for less notational clutter. **NB:** *P*_0_ **is what we actually observe experimentally**.

### A.3 Requirement of no additional conjugation events

One problem with this approach is that it ignores the possibility that another conjugation event may have occurred in this 5 minutes. The probability of 0 plasmids at *t* = 5 is implicitly the probability of (degradation within 5 minutes) AND (no additional conjugation within 5 minutes). The probability of no additional conjugation is given by 1 minus the cdf of the exponential distribution with rate *k*_*c*_ where *k*_*c*_ is the rate at which plasmids are conjugated into the cell: 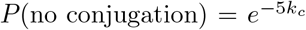. (If e.g. *k*_*c*_ = 0.07 this probability becomes 0.70). We assume that degradation and conjugation are independent, so that

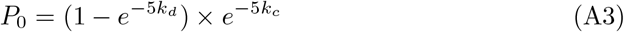

which yields

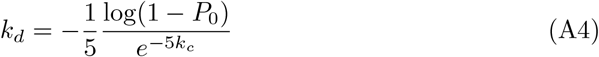

Since 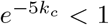, this will lead to a larger estimated value for *k*_*d*_ than with the naive approach. For instance, if *k*_*c*_ = 0.07, then 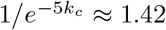.

### A.4 Possibility of additional conjugation and degradation events

In reality, there are numerous “trajectories” that could lead to observation of 0 plasmids at *t* = 5 starting from one plasmid at *t* = 0. For instance the original plasmid could have been degraded AND any number of additional conjugation and degradation events could have occurred as long as the number of additional degradation events matches the number of additional conjugation events. So to correctly compute *P*_0_, we need to sum over all trajectories yielding 0 plasmids after 5 minutes. Let *T* denote plasmid copy number trajectories. Then

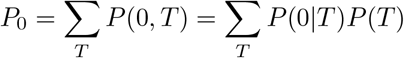

But *P* (0|*T*) = 1 for “valid” trajectories and 0 otherwise, so

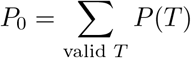

Valid trajectories are trajectories where (the initial plasmid is degraded) AND (any subsequently conjugated plasmids are degraded). Let *d*_*i*_ represent degradation of the original plasmid, and *c* and *d* represent conjugation and degradation of any subsequent plasmids.

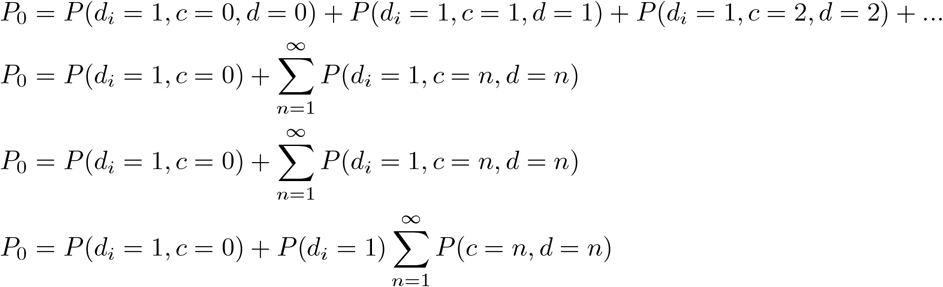

where we assumed that degradation *d*_*i*_ of the initial plasmid is statistically independent of conjugation and degradation of any subsequent plasmids. If we further also assume that any future conjugation/degradation paired events are statistically independent from one another then we get

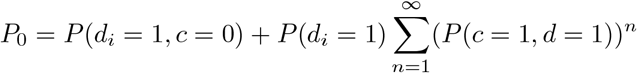

In equation A1 we found that 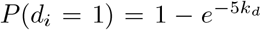 and in equation A3 we found that 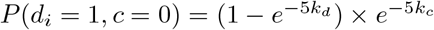. So thus far we have

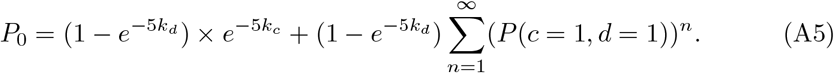

#### A.4.1 Calculation of *P*_*GL*_ = *P* (*c* = 1, *d* = 1)

What is the probability of one conjugation and one degradation event occuring within the next 5 minutes? Let’ s call this *P*_*GL*_ (for “gain and loss”). It is the (probability of conjugation at time *t*^*′*^) times (the probability of no additional conjugation events within 5 − *t*^*′*^) times (the probability of degradation within 5 − *t*^*′*^) summed over all *t*^*′*^ from 0 to 5 minutes. (Strictly speaking the first parenthesis should say: probability of conjugation between *t*^*′*^ and *t*^*′*^ + *dt*^*′*^). Or

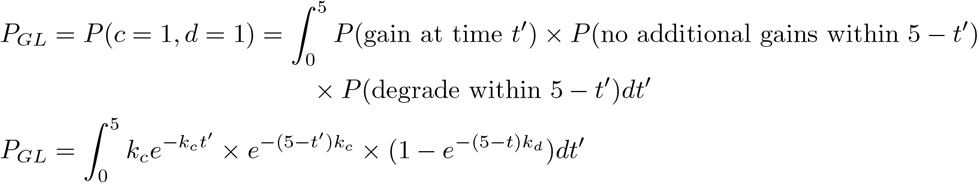

where used the fact that the first term in the integral (gain at time *t*^*′*^) is given by the exponential distribution with rate *k*_*c*_, the second term (no additional gain within 5 − *t*^*′*^) is 1 minus the cdf of the exponential distribution with rate *k*_*c*_, and the third term (degrade within 5 − *t*^*′*^) is the cdf of the exponential distribution with rate *k*_*d*_ over the interval 5 − *t*^*′*^. After some algebra we find that

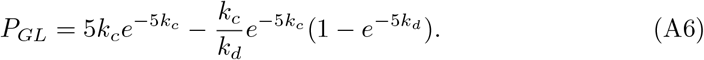

The first term can be straightforwardly interpreted as the probably that exactly one conjugation occurs during the 5 minutes. The second term is slightly less transparent but should reflect the probability that the plasmid does not get degraded in the time after conjugation. Looking at some limits, if 5*k*_*c*_ *>>* 1 (fast conjugation) the probability goes to zero since the chance of only one conjugation is small. If *k*_*d*_ *>>* 1 (fast degradation) then the second term goes to zero and the probability reduces to the first term i.e. the probability of having one conjugation event. Makes sense.

#### A.4.2 Putting it together

Plugging these results back into equation A5 we obtain

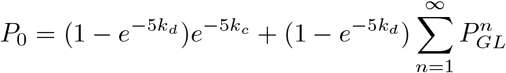

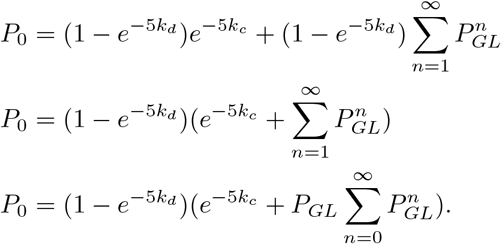

Finally, using the formula for the sum of geometric series and the fact that *P*_*GL*_ < 1 we find

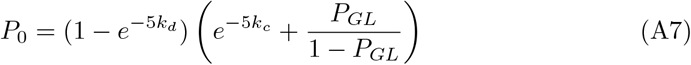

where

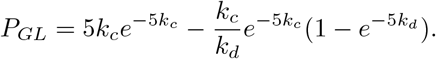

This will need to be solved numerically for *k*_*d*_, assuming that *k*_*c*_ is known from non-CRISPR-targeting experiments. Note that if *P*_*GL*_ *<<* 1 this reduces to equation A3. As an example, if *k*_*c*_ = 0.07 and *k*_*d*_ = 0.2, we obtain *P*_*GL*_ = 0.09. Which seems reasonable.

### A.5 Requiring two consecutive plasmid-free frames

We might choose to require that two consecutive plasmid-free frames are required to call a clearance event, in order to cut down on false negatives (or false positive for clearance, depending on how you look at it). In this case *P*_0_ becomes the probability of 0 plasmids at *t* = 5 AND *t* = 10 given that one plasmid was observed at *t* = 0. Or

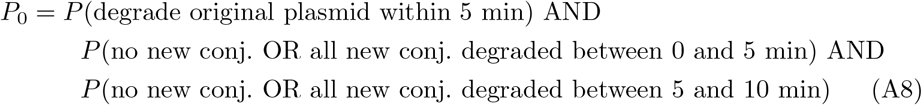

In other words we just need to add another factor of the second term in equation A7:

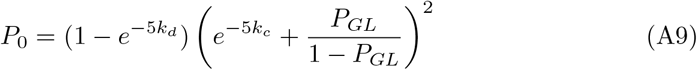

### A.6 Correcting for false negatives

So far we have mostly avoided considering the possibility of false negatives for plasmid presence. We can estimate the probability of false negatives (no plasmid is detected when plasmid is present) from experiments with non-targeted plasmids, where we assume that all plasmid loss in this case is due to false negatives. (This may not be 100% true if e.g. cell division occurs in a newly-conjugated cell before the plasmid has a chance to replicate, but it’ s probably still a good-enough assumption since few divisions will occur within 5 minutes).

What we have been calling *P*_0_ - the probability of zero plasmids at *t* = 5 given one plasmid as *t* = 0 - is really the probability of *observing* zero plasmids which may not be the true probability. We can write the total probability of observing zero plasmids as a sum of the probabilities of observing zero plasmids given that true number of plasmids is either zero or greater than zero:

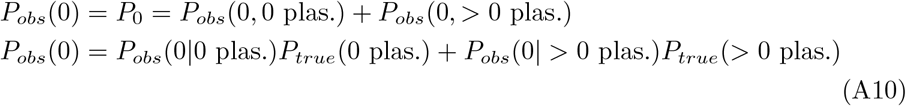

We assume that dot detection parameters have been tuned such that the probability of false positives is negligable and hence *P*_*obs*_(0|0 plas.) = 1. We define the probability of false negatives *P*_*FN*_ = *P*_*obs*_(0| *>* 0 plas.) and use the fact that there must be either zero or greater than zero plasmids so that *P*_*true*_(0) + *P*_*true*_(*>* 0) = 1 to obtain

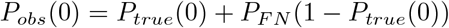

which after rearranging yields

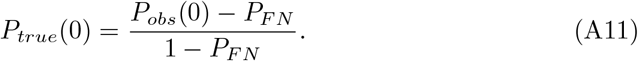

This seems reasonable because we essentially subtract the probability of false negatives and then normalize to the size of the interval remaining after subtracting *P*_*FN*_ so that *P*_*true*_ again lies between 0 and 1.

To use this expression we simply substitute *P*_*true*_ into e.g. equations A7 or A9, using the appropriate estimated *P*_*FN*_. For context we typically see *P*_*FN*_ values around 0.07 when two plasmid-free frames are required and 0.17 when one plasmid-free frame is required.

### A.7 Limitations

This all becomes much harder if plasmid replication is included. This assumption should be revisited; however, it is not unreasonable to assume that a 60 kb plasmid takes longer than 5 minutes to replicate. Also, RP4 carries the entry exclusion gene *trbK* which prevents conjugation into cells already harboring RP4. In the preceding analysis, we asssume that TrbK protein concentration does not build up sufficiently in 5 minutes to confer the entry exclusion phenotype.

## Appendix B Agent-based models of population dynamics

### B.1 Overview

As mentioned in the main text, the agent-based simulations consist of two partially-independent modules controlling biochemistry (plasmid conjugation, replication, and clearance) and physics (cell positions, forces, and movements) respectively (Fig. S3). The main conceptual challenge lay in reconciling the fixed time steps Δ*t* (1.25 min in our case) of the physics module with the variable time steps of the Gillespie algorithm (briefly explained in next paragraph) used in the biochemistry module. Moreover, the modules cannot simply be run independently in parallel because biochemical events such as conjugation depend on a cell’ s neighbors at a given time point, and cell neighbors are governed by the physics module and can change every Δ*t*. The solution we adopted was to run the biochemistry module largely in parallel but to update conjugation rates every Δ*t*. This means that at each simulation step, for each cell, Gillespie steps (Fig. S3B) are taken starting from *t* = *t*_*sim*_ until a step occurs later than *t*_*sim*_ + Δ*t*. This final Gillespie step is discarded, and the Gillespie simulation for that particular cell is restarted at the next simulation step starting from *t*_*sim*_ + Δ*t* with appropriately updated rates. By “simulation step” we mean an iteration encompassing the two modules as shown in Fig. S3A. By “Gillespie step” we mean one step of the Gillespie algorithm simulating changes in plasmid copy number within a given cell.

The Gillespie, or stochastic simulation, algorithm is a method to simulate stochastic biochemical trajectories based on reaction rates. It has been amply explained elsewhere, but very briefly, for each Gillespie step, the sum of rates *k*_*tot*_ for all reactions is calculated. For instance, for transconjugants, this would be the sum of plasmid replication and plasmid clearance rates (Fig S3B). The time interval *dt* until the next event happens is then drawn from an exponential distribution with mean 1*/k*_*tot*_. The identity of the next event is randomly chosen weighted by the relative rates of the possible events. The number of molecular species are then updated according to the reaction chosen, and reaction rates are re-calculated. The next timestep *dt* is chosen based on the updated rates, and so on. In our case, Gillespie steps are taken until the last step oversteps *t*_*sim*_ + Δ*t* as described above.

### B.2 Simplifications

Our overarching aim was to find the simplest set of rules that could explain the experimental data and so certain simplifications were made while creating the simulation:

- Cell growth rates were independent of cell classes and Cascade expression levels, Justification: the differences are single digit percents and should not significantly affect population statistics over 75 minutes, which is less than two generations at *≈* 1.0 h^−1^ growth rates.
- Cascade expression (or more accurately, Cascade concentration) remained the same over the course of the simulation for cells and their respective offspring. This allowed us to avoid explicitly simulating (stochastic) gene expression, which is a nontrivial project in itself. Justification: Cell were subcultured for three hours in identical growth media prior to the experiment and expression does not change significantly over less than two generations.
- We chose to limit simulation lengths to 75 minutes because in our experiments this was generally the interval within which most of the actual population dynamics occurred.
- We deemed simulating the effects of ParE toxin to be beyond the scope of this work and therefore focused on Δ*parE* variants in simulations.

### B.3 “Burn-in” of cell positions

The physics module models cell as rigid rods, whereas real cells are not perfectly rigid. We found that taking the literal positions and cell lengths from experimental data often caused the physics module to crash since cells overlapped by too much. To sidestep this issue, prior to running simulations, we performed a “burn-in” run where cell lengths were first reduced by 50%, and then grown using the physics module until reaching their original lengths. During this burn-in phase, no conjugation or other changes in internal cell state took place; cell growth was the only aspect allowed to change. This also explains why some cells are positioned slightly differently in Fig. 6A vs. Fig. 6C.

### B.4 Plasmid replication

Control of plasmid replication and copy number is a field of research in itself; here our objective was to create a simple empirical model to describe the observed plasmid copy numbers in the absence of CRISPR interference. We found that the average plasmid copy number increases linearly with cell length for 0-target plasmids. We identified the observed average copy number as the “goal” plasmid copy number *n*_*goal*_ (Supplementary Fig. S3B), which in turn depends on cell length. We reasoned that, upon conjugation into a new cell, plasmids would initially replicate as fast as possible, but reduce their replication rate as plasmid copy number approached *n*_*goal*_. We modeled this by postulating that the replication rate per plasmid decreases linearly from *k*_*rep,max*_ when *n*_*plas*_ = 1 to zero when *n*_*plas*_ = *n*_*goal*_, where *k*_*rep,max*_ is the maximal replication rate, yielding the following expression for replication rate:

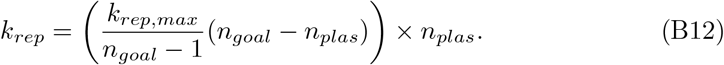

### B.5 Data included in simulations

We typically imaged 10 microfluidic traps (positions) per donor-recipient pair per experiment. In Fig. 6ABC, the initial positions and cell classes are shown for a single trap. However the population dynamics (experimental and simulated) in Fig. 6DE are based on averaging over all 10 traps for that particular construct. For Fig. 6F, we used 5 positions as training data to determine the optimized value of *k*_*clear*_, and 5 positions to test this value. Thus Fig. 6F reflects the 5 “test” positions.

### B.6 Approximate Bayesian Computation

ABC entails comparison of a chosen summary statistic between simulation and experiment. We chose to use the fraction *T/*(*T* + *R*) of recipients that have been “converted” to transconjugants as our summary statistic. We experimented with different summary statistics but found that e.g. the donor fraction was relatively uninformative because it only slightly varied in a manner that largely reflected random fluctuations in trap loading. *T/*(*T* + *R*) on the other hand seemed to more accurately reflect the dynamics of conjugation.

We required that “accepted” simulation trajectories have a *T/*(*T* + *R*) value with a mean absolute distance from the experimental *T/*(*T* + *R*) trajectory less than 0.03.

